# Generating Interacting Protein Sequences using Domain-to-Domain Translation

**DOI:** 10.1101/2022.05.30.494026

**Authors:** Barthelemy Meynard-Piganeau, Caterina Fabbri, Martin Weigt, Andrea Pagnani, Christoph Feinauer

**Affiliations:** Department of Computing Sciences, Bocconi Institute for Data Science and Analytics (BIDSA), Bocconi University, Milan, Italy; Politecnico di Torino, Corso Duca degli Abruzzi, 24, I-10129, Torino, Italy; Sorbonne Université, CNRS,, Institut de Biologie Paris Seine, Biologie Computationnelle et Quantitative LCQB,, F-75005, Paris, France; Italian Institute for Genomic Medicine, IRCCS Candiolo, SP-142, Candiolo, Italy; INFN, Sezione di Torino, Torino, Via Pietro Giuria, 1 10125 Torino Italy

## Abstract

**Motivation:** Being able to artificially design novel proteins of desired function is pivotal in many biological and biomedical applications. Generative statistical modeling has recently emerged as a new paradigm for designing amino acid sequences, including in particular models and embedding methods borrowed from Natural Language Processing (NLP). However, most approaches target single proteins or protein domains, and do not take into account any functional specificity or interaction with the context. To extend beyond current computational strategies, we develop a method for generating protein domain sequences intended to interact with another protein domain. Using data from natural multi-domain proteins, we cast the problem as a translation problem from a given interactor domain to the new domain to be generated, i.e. we generate artificial partner sequences conditional on an input sequence.

**Results:** Evaluating our model’s quality using diverse metrics, in part related to distinct biological questions, we show that our method outperforms state-of-the-art shallow auto-regressive strategies. We also explore the possibility of fine-tuning pre-trained large language models for the same task and of using Alphafold 2 for assessing the quality of sampled sequences.

## 1 Introduction

Generating novel protein sequences with desired properties is one of the key challenges of computational biology. It is likely that machine learning methods will play an important role in this task, being already used for the generation of new enzymes, biological sensors and drug molecules [35]. A promising approach is to leverage deep generative models, which use neural networks for learning probability distributions from known, naturally occurring protein sequences [2, 14, 20, 27, 30]. Apart from other uses, like the prediction of mutational effects [28], these models can be used for protein design by selecting high-probability sequences (possibly under constraints) from the learned distribution.

Naturally occurring protein sequences are often comprised of several domains, and domains can be classified into different families [1]. Models that work on the domain level usually use as training data a single multiple sequence alignment (MSA) [8], containing sequences from the same domain family after aligning them, and make the assumption that each sequence is constrained by the same fitness landscape. This modeling paradigm neglects the dependence of the sequence constraints on the specific context corresponding to each organism, including other proteins interacting with the sequence or other domains on the same protein. Together with the fact that most of the crystallographic structures deposited in the PDB database [5] are resolved only at the single domain level [37], this poses interesting questions about the limitations of current approaches, for example when predicting the relative orientation of multi-domain proteins [35]. Another field where this issue arises is immunology, where monoclonal antibody experiments are typically performed on mouse models and only later tested in humans. This is related to the so-called humanization problem, i.e. how to graft a promising variable receptor region (CDR) from a murine to a human context [6].

Known families of interacting domains can be organized in a paired multiple sequence alignment (pMSA), where the aligned interaction partners are concatenated [24]. Given the evolutionary pressure for maintaining functional interactions between proteins, amino acid substitutions at interaction surfaces are not independent between the interaction partners. The interacting sequence therefore can be used as additional information when generating a novel sequence. The current work addresses the task of generating domain sequences given an interacting domain sequence. Given that this task is similar to translation tasks in natural language processing (NLP), we explore the use of Transformers in this context. While there is some recent work using Transformers for translating between protein sequences [34] for specific applications, there is, to the best of our knowledge, no systematic exploration of this idea on the level of protein domain families on a diverse dataset. We explore different architectural choices, regularization schemes and compare our results with a recently published shallow auto-regressive method [32], which we use as a baseline. We also compare on a smaller scale to fine-tuned large protein language models, using RITA [15], and explore how structural predictions from Alphafold 2 correlate with our results.

The general idea of this work is summarized in Fig. 1. Consider a protein with at least two interacting domains, where interaction is defined as having a pair of amino acids at a distance less than 8 angstrom. We then search a database of proteins for other sequences where these domains co-occur in the same protein and assemble the pMSA and use it for training the Transformer to translate from one domain to the other. The decoder being a causal language model, we can efficiently calculate the probability of a target sequence give the input sequence. This probability enables us to evaluate the compatibility of domains, which can be used for matching a domain to an interacting partner among several possible partners. The model is generative in that it can be used for generating a novel target sequence given the input sequence. Given a context, we can generate a new “translation” or target sequence and evaluate the new de novo proteins.

**Figure 1:**
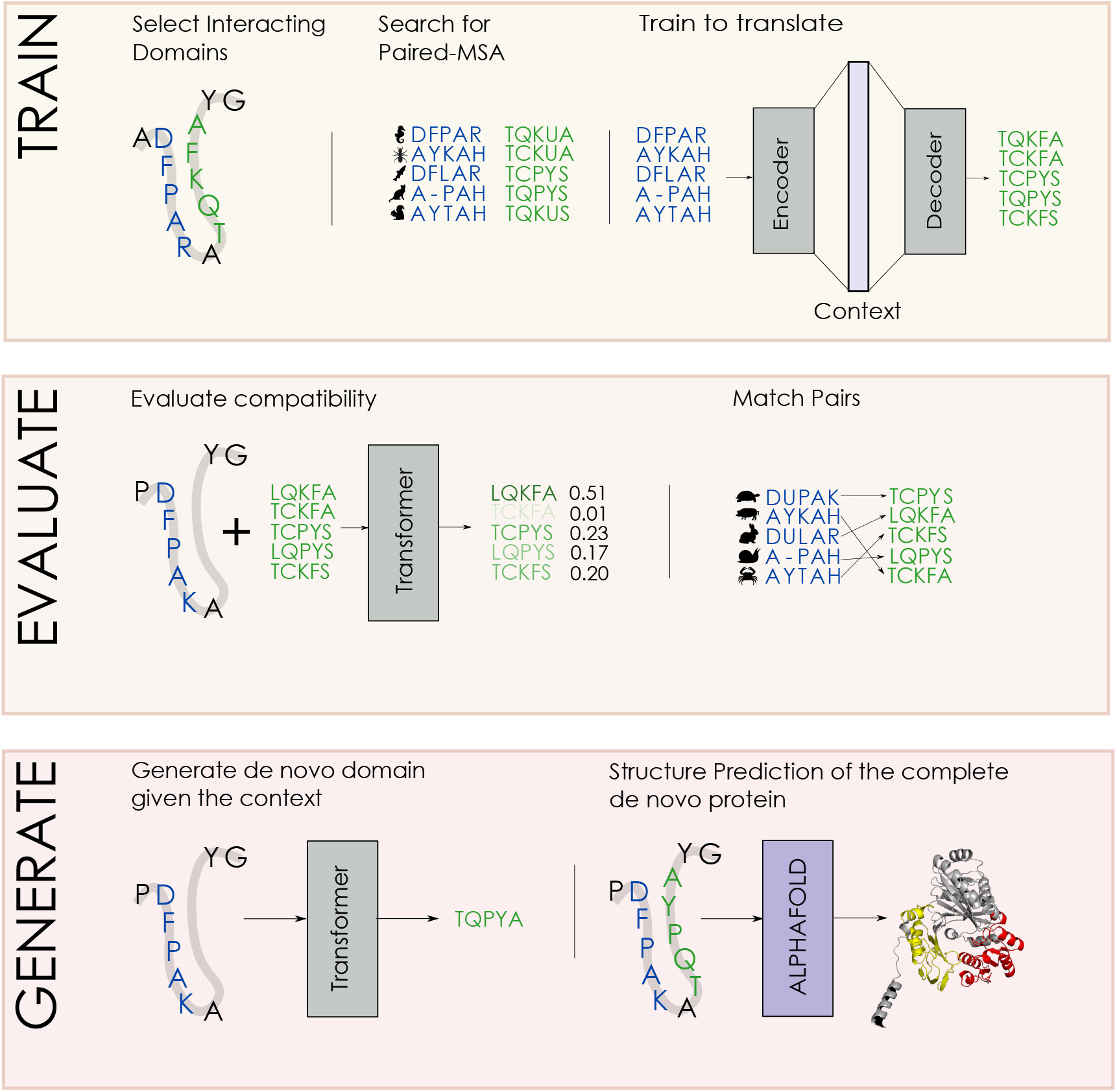
Summary of the work presented in the paper. In the first box (Train) we extract interacting domains from known structures. We then build a pMSA based of homologous sequences of these domains and train the Transformer to translate between them. In the second box (Evaluate), we use the probabilities of the trained Transformers to match source and target domains and assess the resulting accuracy. FIn the third box (Generate), we sample novel target domains and use it for replacing the original target domain. We the use Alphafold to predict the structure of the modified sequence and analyze the difference to the original structure

## 2 Related Literature

Generative modelling for protein design has a wide range of applications and a considerable number of different models have been proposed in literature, recently especially deep neural network models [35]. These include autoregressive models based on convolutional architectures [30], generative adversarial networks [27], variational autoencoders [14], LSTM based architectures [2] and self-attention based architectures [20]. The latter work allows for sequence generation conditioned on tags corresponding to molecular function or taxonomic information. Similar to results in NLP, scaling protein language models to very large sizes seems promising for protein sequences [15].

Transformer-based architectures [33], which we use in the present work for sequence-to-sequence prediction, have also been used, for example, for creating generic embeddings trained on almost all known protein sequences [29], the prediction of mutational effects [22], protein interaction prediction and protein family classification [25], MSA-based language modelling [26], protein contact prediction [36], inverse folding [16, 21] and have been at the core of recent breakthroughs in protein structure prediction [18].

Recently, specific tasks have been cast as sequence-to-sequence translation problems using Transformers, similarly to our approach. This includes for example the generation of drug molecules given a protein sequence the molecule should interact with [12] and the generation of short signal-peptides guiding the secretion of industrial enzymes, given the amino acid sequence of the enzymes [34].

Finally, non-neural network models borrowed from statistical mechanics have been extensively used in the context of sequence generation, for example Generalized Potts Models, a particular form of Markov Random Field [10]. This type of model can be used for generating sequences using MCMC strategies, albeit with a significant computational cost. Relevant approximation strategies are for example the recently introduced auto-regressive (shallow) variants [32], which show a similar performance to Potts models but are computationally more efficient.

## 3 Data and Methods

### 3.1 Dataset

Our data consists of 27 pMSAs containing domain sequence pairs that are part of the same multi domain proteins, taken from [24]. The dataset contains only domain pairs which form a structural contact in at least one resolved PDB structure, making it likely that the two domains co-evolve in order to maintain compatibility. Each dataset is comprised of *M* rows corresponding to *M* sequence pairs, where *M* depends on the dataset and ranges from a few hundred to more than 15.000, see Appendix. Sec A for a summary of the datasets used. The sequences are already aligned using standard bioinformatics tools [11], which means that sequences belonging to the same domain family have the same length. Each row *l* in a dataset represents a pair of domain sequences *B*^*l*^ and *A*^*l*^, which are part of the same protein. Every sequence consists of symbols denoting either one of 20 amino acids or an alignment gap, making the total size of the vocabulary equal to 21.

The first sequence 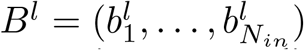 is called the source or input sequence and we denote its length by *N*_*in*_. It is used as an input to predict the second sequence, called the target or output sequence, 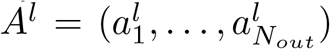, which is of length *N*_*out*_. All source sequences 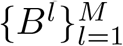 in the pMSA are members of the same domain family, and all target sequences 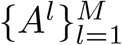 in the pMSA are members of the same domain family. Each dataset was randomly split into a training set (70%) and a validation set (15%). The last 15% were kept for a testing set in order to be able to optimize hyperparameters for every domain, but we did not use it in the experiments shown in this work. Since sequences in an MSA are related to each other due to phylogeny, the validation set might contain sequences that are nearly identical to some sequences in the training set. We therefore further divided the validation set into two parts, one close to the training set and one far from it. This allows us to control for the effects of phylogeny on the performance metrics. This second splitting was made based on the median of the Hamming distance from the training set. The details for this subpartition are found in the Appendix Sec. A.

### 3.2 Performance Metrics

#### Log-Likelihood and Perplexity

An interesting property of autoregressive models, such as the Transformer or arDCA, is that they define a tractable probability distribution over the space of sequences. Contrary to, e.g., Potts Models and other energy based models, we do not have to evaluate a global normalizing constant over the complete space of possible sequences. We can therefore calculate the log-likelihood of a sequence *A* given *B* as

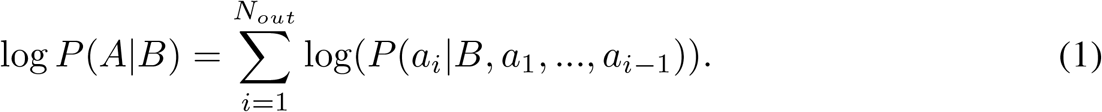

This is related to the cross-entropy, which we use as a loss during training,

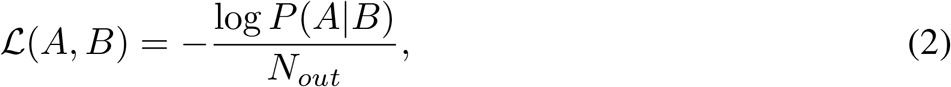

which we average over batches during training.

For assessing one aspect of the quality of our models, we use the closely related perplexity 𝒫𝒫 (*A, B*), which is a common quality metric for protein language models [3], and can be defined as

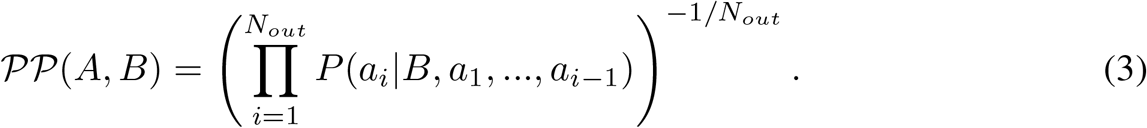

Below we show averages of the perplexity over the training and validation sets and use the notation 𝒫𝒫^*train*^ and 𝒫𝒫^*val*^ for these.

#### Accuracy

While we use the perplexity as one metric for the quality of our model, it is not always easy to interpret: A high perplexity can result from a single wrong prediction with a high level of confidence. We therefore also use the accuracy 𝒜 (*A, B*) for assessing our models. This measure takes the same input as the cross-entropy (the conditional probability for every position) and counts the fraction of times where the true amino acid is the one with the highest probability, leading to

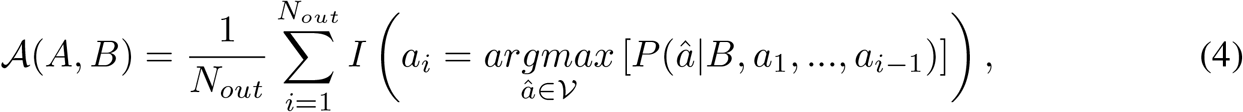

where 𝒱 is the alphabet of symbols and *I* is an indicator function that is 1 if its argument is true, and 0 else. We define 𝒜^*train*^ and 𝒜^*val*^ as the average of the accuracy on the training and validation set.

#### Matching Specificity

We expect the interaction between two domains to affect the probability distribution of the target sequence only marginally, with much of the variability in the distribution being explainable by constraints internal to the target sequence. As a consequence, a good performance in the quality measures defined above might be due to the decoder being a good language model of the target protein, possibly ignoring the input sequence altogether. We therefore also evaluate the specificity of the predicted target sequence given the source sequence.

Specificity is also related to the task of matching pairs of protein sequences, which is an active domain of research in bioinformatics [4, 13, 31]. We implement this task by separating the source and target sequences in the validation pMSA, resulting in two separate MSAs with the same number of rows, one containing the source sequences and one the target sequences. We then shuffle the rows in the target MSA randomly and attempt to use our models to find the permutation of the target sequences that matches the original order. In order to create a matching based on a model, we calculate the log-likelihood of every combination of source and target sequences in the shuffled validation set and create a matching between source and target sequences based on the Hungarian algorithm [19].

We then use the ratio of correctly matched pairs as an additional metric for the performance of our model, formally defining it as

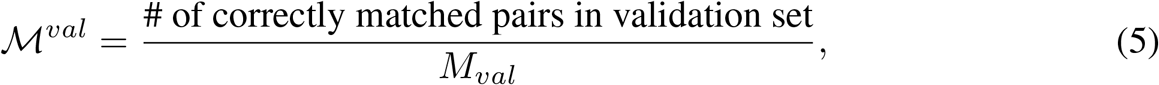

where *M*_*val*_ the size of the validation set. Note that the difficulty of this task increases with the size of the validation set, since the expected ratio of correctly matched pairs using a random matching is 1*/M*_*val*_.

### 3.3 Transformers and Baselines

We used two Transformer models with different sizes, calling one the *shallow* and one the *large* model. The shallow Transformer consists of 2 layers with a single attention head, has an embedding dimension of *d*_*model*_ = 55 and a forward dimension of *d*_*ff*_ = 2048. The large Transformer consists of 3 layers and has an embedding dimension of *d*_*model*_ = 105 with the same forward dimension and number of heads as the shallow transformer. Further details on their architectures can be found in Appendix Sec. B.1. We compare their performance to the recently introduced shallow auto-regressive model called arDCA [32] and a fine-tuned version of RITA L [15]. While details on these methods can be found in Appendix Sec. B.2, we note here that RITA was pre-trained on a large corpus of unaligned, full-length sequences, which is a different setting from the pMSAs that we use for the Transformers. We therefore evaluated RITA only on unaligned, full-length sequences. For arDCA, which we train from scratch on pMSAs, there is no such mismatch and we can use it on the same pMSAs as the Transformers. For the datasets used in this work the training time of the Transformer models ranges from less than an hour to about 1.5 days for the large Transformer on the largest dataset. Training was done using a single Nvidia V100 GPU. When using entropic regularization, which we will introduce in a later section, the training time increases significantly. We provide a table with training times in Appendix Sec. B.4.

### 3.4 Entropic Regularization

When experimenting with the large Transformer, we observed strong overfitting of the perplexity, especially when trained on smaller datasets. While this could be expected, we found that the matching performance was not following the same trend: While the perplexity started to degrade at some point during training, which is indicative of overfitting, the accuracy and the matching performance were still increasing, see Appendix Sec. C.1. While the shallow Transformer is less prone to overfitting, most likely due to its limited capacity, we found it necessary to introduce regularization for the large Transformer. We experimented with dropout and weight decay with limited success. While both schemes prevent overfitting in terms of the perplexity, the matching performance and the accuracy dropped significantly. We show this effect in Appendix Sec. C.1 for different training set sizes and regularization settings.

In order to find models with a good performance on perplexity, matching and accuracy at the same time, we explored other regularization approaches.

In this section, we present an approach based on entropic regularization, where we enforce the probability of a target sequence *A* given a source sequence *B* to be similar to other sequences sampled from the model conditioned on *B*. This encourages the model to give similar weights to different possible interaction partners, even if there is only a single one present in the training set.

We therefore add a regularization term 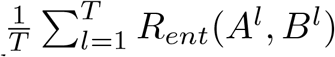 to the loss, where *l* indexes the input sequence *B*^*l*^ and the target sequence *A*^*l*^ in the batch and *T* is the batch size. We sample *S* different target sequences for *B*^*l*^ from the model. We denote the *k*^*th*^ sampled sequence conditioned on *B*^*l*^ as *A*^*l,k*^. We sample using a Gumbel-Softmax distribution [17], which enables back-propagation through the sampling step. For computational efficiency, we sample every amino acid in *A*^*l,k*^ conditioning on the preceding amino acids of the true *A*^*l*^. Then we evaluate the log-likelihoods *R*^*l,k*^ of the target sequence *A*^*l,k*^ given *B*^*l*^ and the log-likelihood of the true pair *R*^*l*^,

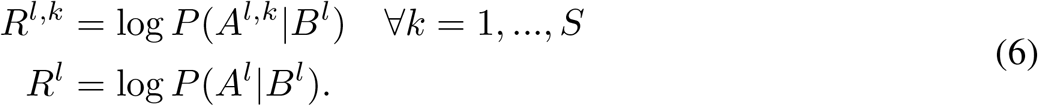

We then use these quantities as the input for a log-softmax operation, resulting in

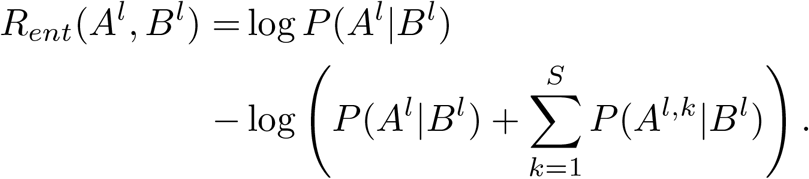

This term is multiplied by a factor *α >* 0 to regulate its strength and *added* to the loss function, meaning that we aim to minimize it. This enforces similar probabilities for the true target sequence *A*^*l*^ and the sampled target sequences *A*^*l,k*^, conditioned on *B*^*l*^. A diagram summarizing the regularization approach can be found in Fig. 2. A closer look reveals that it is a form of entropic regularization, maximizing the conditional Rényi entropy of order 2, see Appendix Sec. C.2.

**Figure 2:**
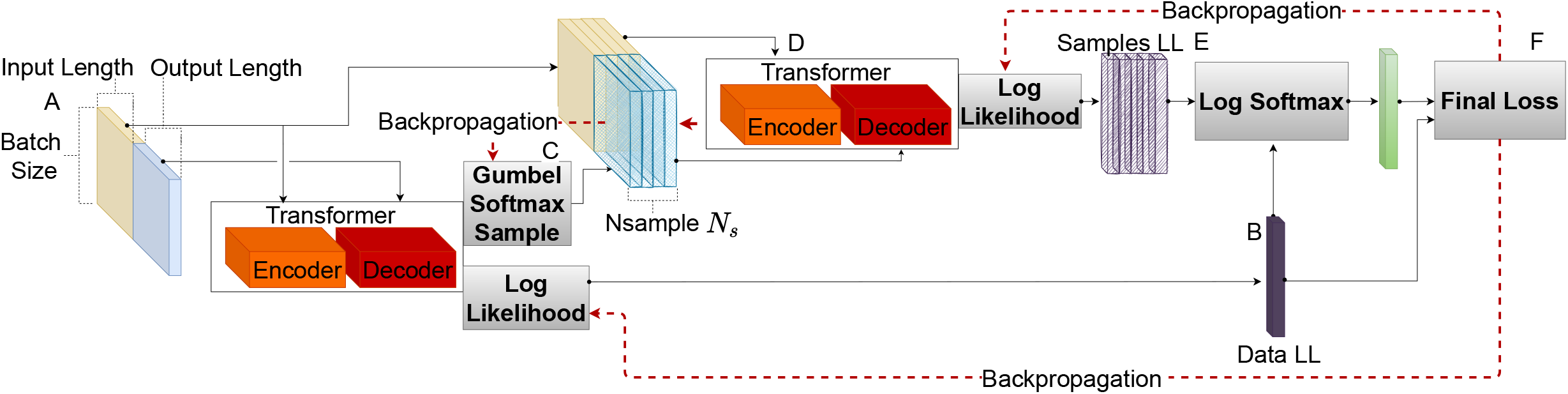
Diagram explaining training with entropic regularization: (A) corresponds to the training batch, with yellow being the input protein sequences and blue the output protein sequences; At (B), the batch is sent to the Transformer and the loglikelihood *Data LL* is computed; At (C), *N*_*s*_ output protein sequences are sampled from the Transformer for every input protein sequence using the Gumbel softmax operation; At (D) we evaluate the loglikelihood of the sequences sampled at (C) and call it *Samples LL*; At (E) we measure how well the loglikelihood separates the training sequences from the sampled sequences using a logarithmic softmax, creating an additional loss term; At (F), the losses calculated at (E) and (B) are combined.

## 4 Results

### 4.1 Performance Gain From Context Sequence

We first tested whether the input sequences had any effect on the perplexity of the target sequence. As already mentioned before, this is not self-evident, since the Transformer decoder itself could be a good model for the target sequence distribution without taking the input into account. We therefore trained two shallow Transformer models, one with the the normal training set and one where we randomly shuffled the pairing between input and output sequences. We then evaluated the models on the normal validation sets, without shuffling. We expect that if the model trained on the normal training set exploits the information in the inputs when predicting the output, it should show a considerably lower perplexity than the model trained on a shuffled dataset.

We show the results of these experiments in Fig. 3. As can be seen, the models trained on the normal dataset have a significantly lower perplexity than the models trained on a shuffled dataset. This corroborates and quantifies the idea that domain sequences that appear in the context of a second domain contain information that can be used for modelling the constraints on the sequence of the second domain. We note that the difference in the logarithm of the perplexity, which is equivalent to the the cross entropy, can be seen as an rough estimate of the mutual information between the output and the input. When the input sequence is randomly chosen, there is no correlation between the input and the output, and the corresponding probability can be seen as the marginal probability of the output sequence. We can therefore write

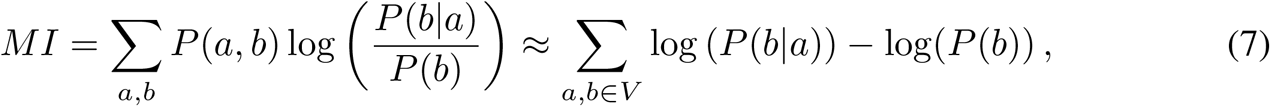

**Figure 3:**
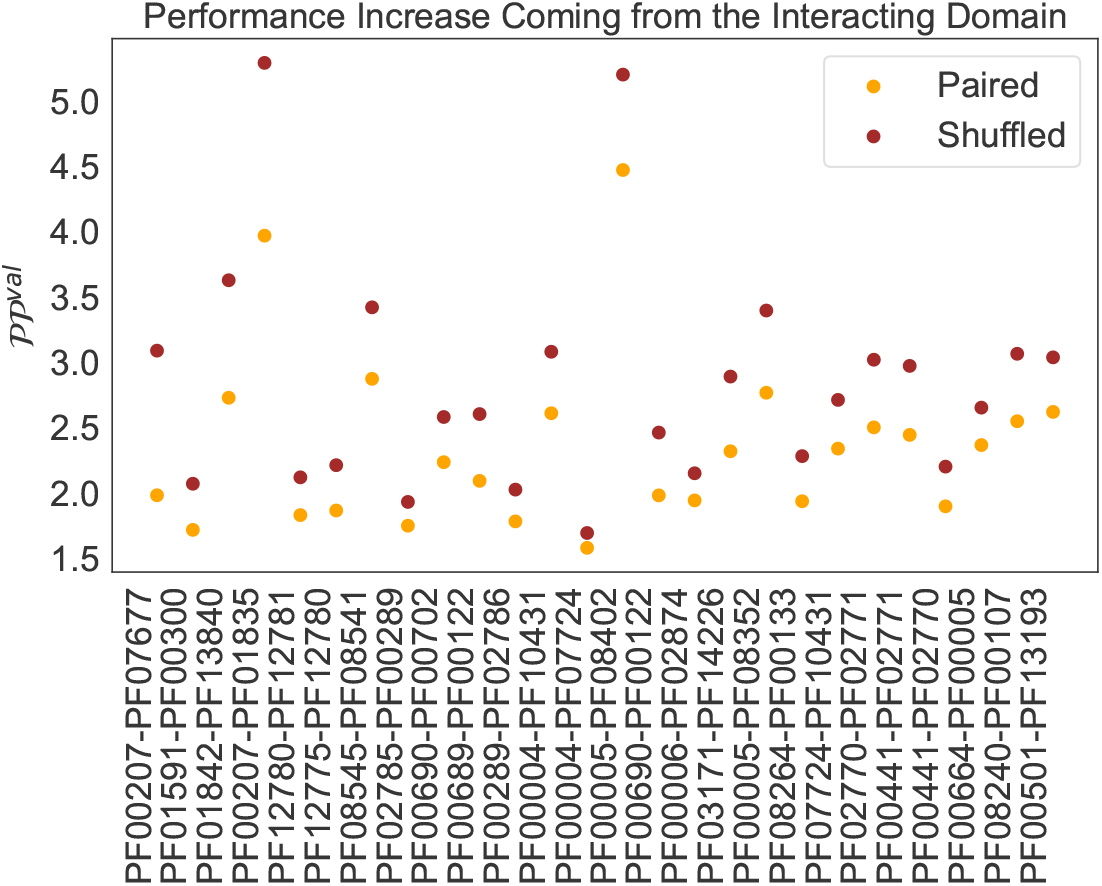
Performance [Lower Better] increase when taking domain sequence in context into account. We plot the Perplexity 𝒫𝒫^*val*^ for target sequences on the validation set, once for a shallow Transformer trained with the true pairings (Paired) and once for a shallow Transformer trained with shuffled pairs in the training set.

where *a* and *b* are sequences in the validation set *V*.

### 4.2 Results on Performance Metrics of the Shallow Transformer

We next compared the shallow Transformer models to the arDCA baseline. Shallow Transformers outperform arDCA on nearly all datasets above a certain training set size in terms of perplexity, accuracy and matching with a large margin, as can be seen in Fig. 4. We note here that the Transformer models (both shallow and large) have *less* parameters than arDCA for every family size we tried: The number of parameters in the Transformer models is independent of the length of the input and target sequences, while the number of parameters in the arDCA models scales quadratically with the concatenated input length.

**Figure 4:**
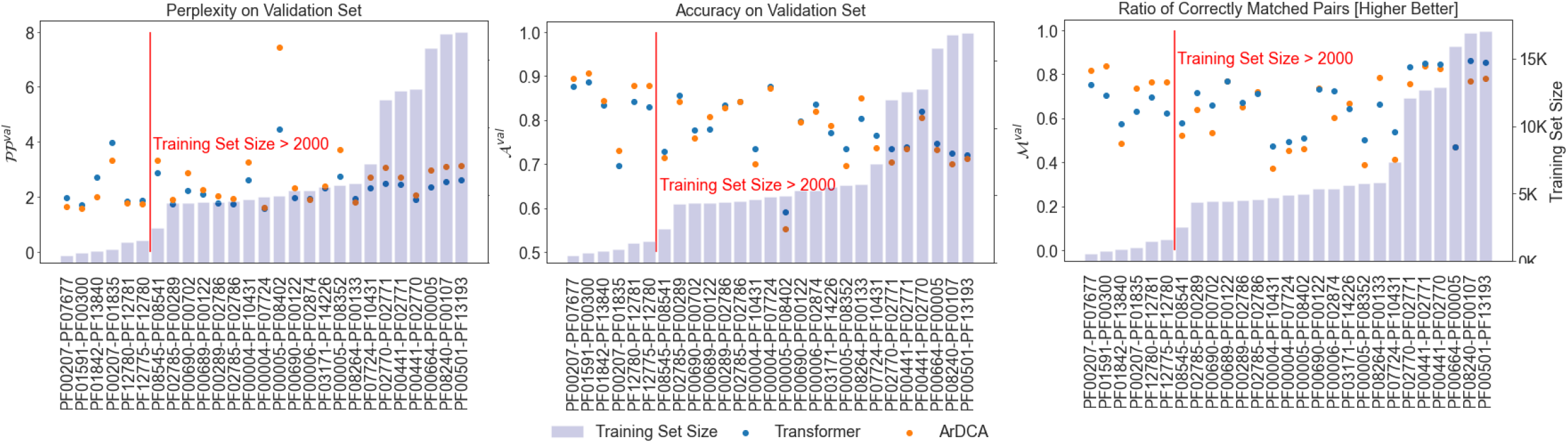
Perplexity 𝒫𝒫^*val*^, accuracy 𝒜^*val*^ and the matching performance ℳ^*val*^ for shallow Transformers and arDCA on validation set. The families are ordered by training set size.

The best performance is achieved for the families with the largest training sets, indicating that the performance of the Transformer might further increase with increasing training set size. We repeated the calculation of the matching performance on subsampled versions of the validation set in order to estimate the dependence on the size of the validation set, see Appendix Sec. F.

We also considered the possibility of fine-tuning a large language model trained on protein sequences for our task. To this end we fine-tuned RITA L [15], a 680M parameters model trained for predicting the next amino acid in a sequence. Given that RITA is trained on full-length unaligned sequences we fine-tune also on full-length unaligned sequences, comparing the metrics only on match positions as predicted by the Pfam HMM of the corresponding domain family (excluding gaps and inserts). The details of the fine-tuning can be found in Appendix.Sec. B.3 The results are comparable with our Transformer models, see Fig. 5, with the Transformer having a slightly higher accuracy. We note that RITA models are trained on Uniref100 and we suspect that most of the sequences in our validation set are in the training set of RITA, so this comparison is likely biased in favor of RITA.

**Figure 5:**
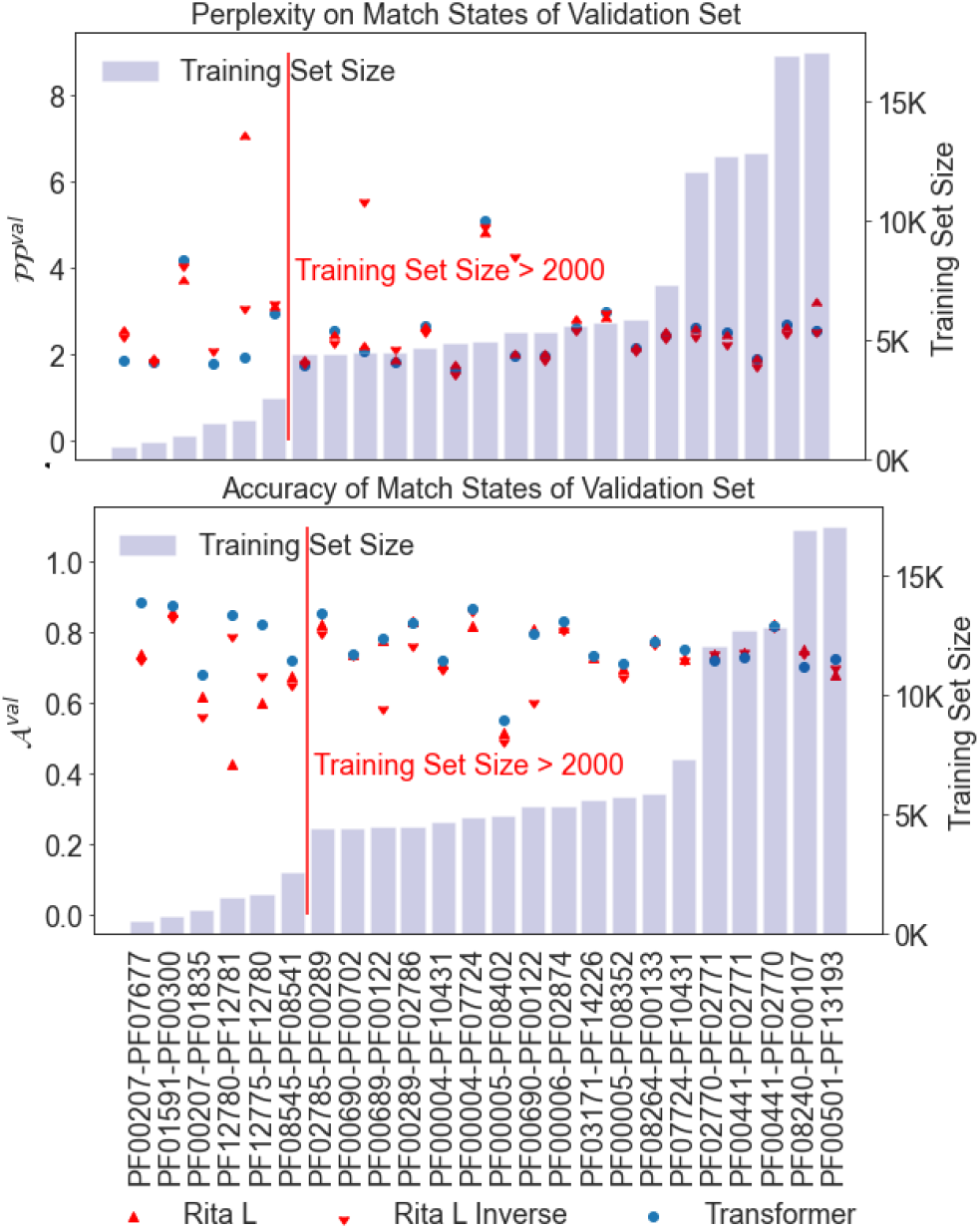
Perplexity 𝒫𝒫^*val*^, accuracy 𝒜^*val*^ and the matching performance ℳ^*val*^ for shallow Transformers and fine-tuned Rita L on validation set. The families are ordered by training set size. Rita L Inverse refers to RITA when given the inverse sequence as input (RITA is trained on both original and inverse sequences).

### 4.3 Performance of the Entropic regularization

We performed a set of experiments on the 27 datasets in order see if this type of regularization improves the performance. We retrained the large Transformer with and without the entropic regularization. We used *S* = 5 and *α* = 0.7 for the experiments. The results can be seen in Fig. 6, where we plot the performance of the shallow Transformer against the performance of the large Transformer for different regularization schemes and arDCA. The details of the training, models and the details of the performance for every family can be found in the Appendix Sec. C.2.1. The large Transformer outperforms the shallow Transformer in terms of accuracy and matching both with and without regularization, indicating that the large Transformer extracted more useful information from the training set. However, the large Transformer without regularization has a significantly higher perplexity on the validation set, indicating overfitting. Adding the entropic regularization leads to a good performance of the large Transformer in all metrics.

**Figure 6:**
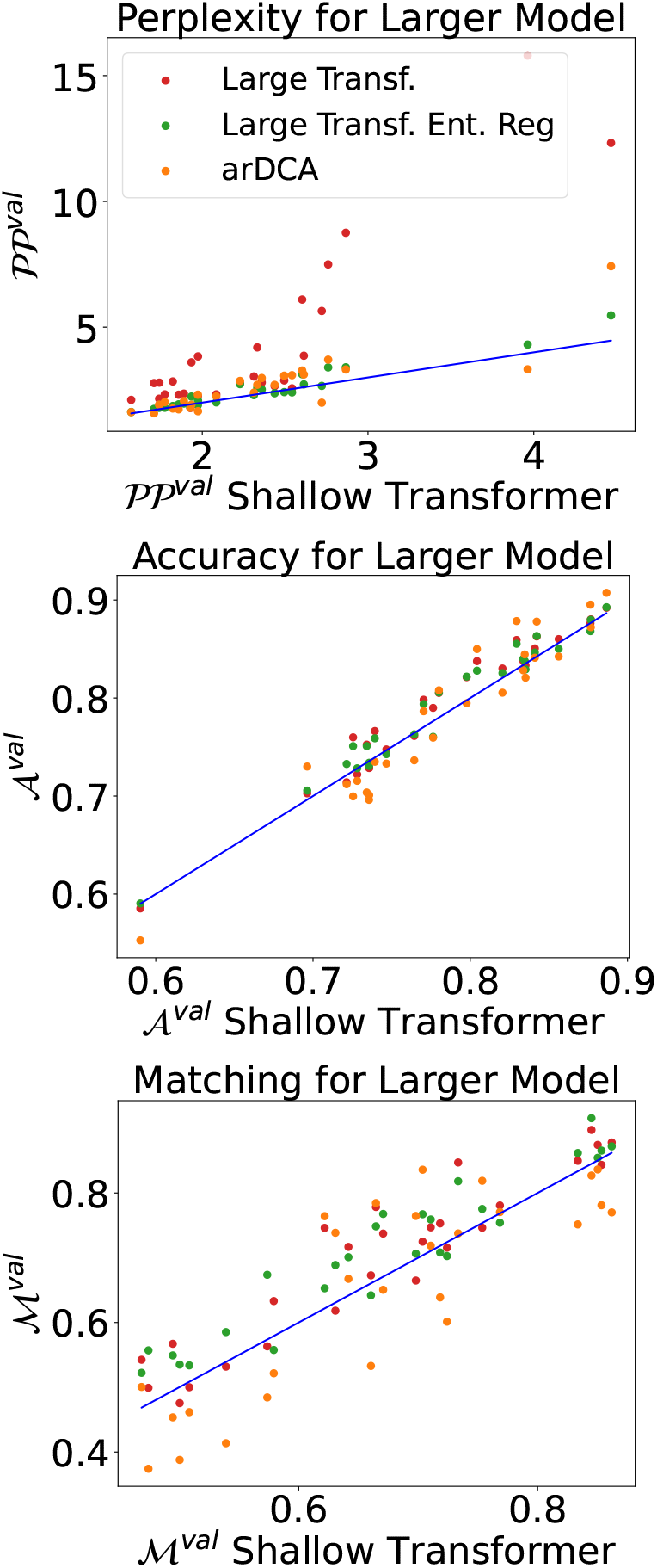
Comparison of the performance of the large Transformer without regularization (red), with entropic regularization (green) and arDCA (orange) and the shallow Transformer. The blue lines have slope 1.

We also performed a systematic comparison of the entropic regularization scheme with standard weight decay, testing different weight decay values for the large Transformer. The details of the experiments and the results for every family can be found in Appendix. Sec. C.3.

### 4.4 Generalization and Phylogeny

One specific characteristic of protein sequences, compared to data in NLP, is the structure of the data. The sequences in our datasets have a phylogenetic bias, visible as clusters of similar sequences in the data, that are simply explained by a close common ancestor. This bias makes a random split unsuitable, since the test set will contain sequences that are very similar to some sequences in the training set’. We therefore evaluate our model on different subsets of the test set, which are selected based on the similarity to the training set.

We show the perplexity on target sequences in the validation set in dependence of the distance from the training set in Fig. 7, where the distance of a sequence to the training set is the smallest Hamming distance from the sequence to any training sequence. Interestingly, it seems that the advantage in performance of Transformer models over arDCA is mostly due to sequences far away from the training set, indicating that Transformers generalize better in regions of sequence space far away from the training set. While Transformers outperform arDCA in matching on sequences both close and far from the training set, the distance from the training set does not seem to systematically influence the performance difference.

**Figure 7:**
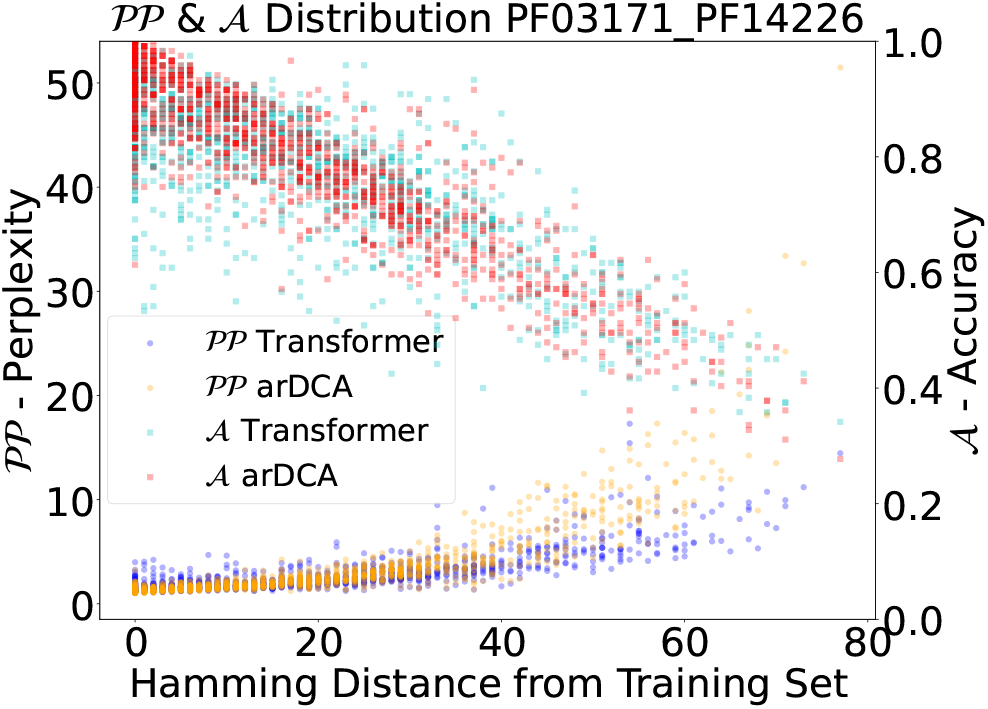
Perplexity (*lower* is better) and accuracy (*higher* is better) for every sequence of the validation set in dependence on the distance of the sequence from the training set. The distance of a sequence to the training set is the Hamming distance to the closest sequence in the training set.

We also verified that this advantage holds for matching. To do so, we split the test set into the half closer and the half further from the training set. When matching the pairs, we look at the performance on these two subdatasets. The details of this results can be found in Appendix Sec. E.1 for the shallow transformer, and at Appendix Sec. C.3 for the Large and regularized Transformer.

### 4.5 Structural

#### Information and Generative Properties

In this section, we show further results on the performance of the shallow Transformer.

We tested whether the target sequence distribution of trained Transformers is integrating structural information. To do so, we explored the correlation of our metrics with scores related to structure prediction when using Alphafold [18]. To this end we selected the protein Q1H158 from the validation set, which contains the Pfam domains PF00289 and PF02785. We then replaced the domain PF00289 with homologous sequences from the validation set, keeping the rest of the Q1H158 sequence unmodified. The resulting sequences contain natural sequences for both domains but in a combination that does not exist in any known protein. We then used Alphafold to predict the structure of the original sequence and the modified sequence, comparing them using the TM-score and the RMSD on the two domains. We found these structural metrics to be well-correlated with the cross-entropy of the resampled PF00289 of the shallow Transformer conditioned on PF02785, see Fig. 8. We stress here that all domain sequences assessed here are natural sequences with presumably a high fitness, which makes it more likely that a higher cross-entropy for a pair is due to a decreased mutual incompatibility, reflected in the structural scores. We present results for more proteins in Appendix Sec. D.1.

**Figure 8:**
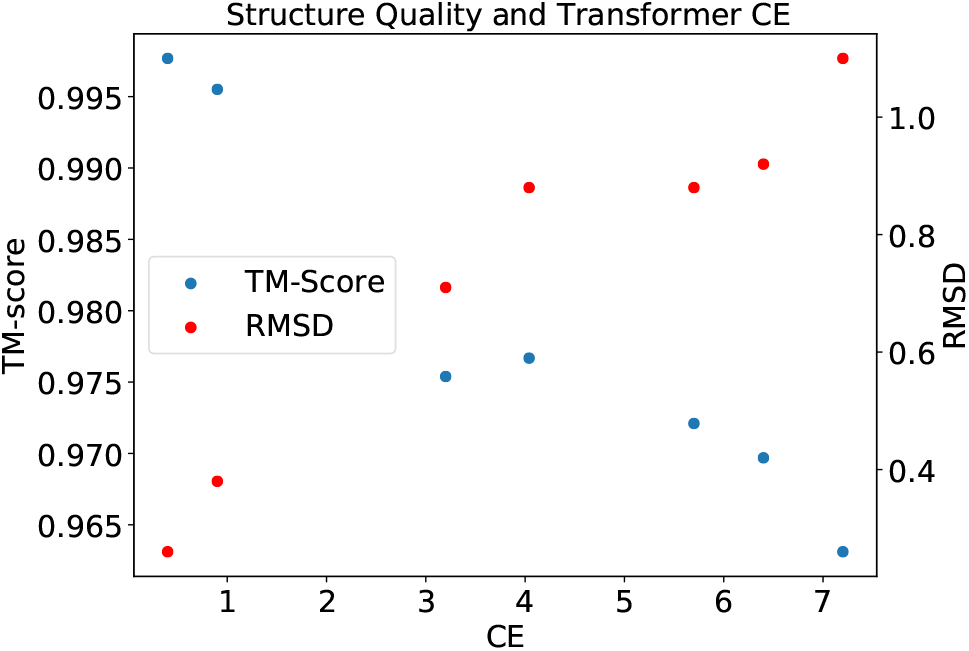
TM-Scores and RMSD values when comparing the Alphafold predicted structures of true sequences with Alphafold predicted structures of sequences where single domains have been replaced with homologous natural sequences. The results are based on Q1H158, which contains domains PF00289 and PF02785, which are in contact in PDB 5ks8. Homologous PF00289 sequences are sampled from the validation set and inserted into the Q1H158 sequence, measuring the change in structural scores and in cross-entropy in the shallow Transformer model (abscissa).

We next assessed the generative power of our Transformer models. To this end, we again used the protein Q1H158 as a test. We sampled novel PF00289 domain sequences conditioned on the PF02785 sequence found in Q1H158 using the shallow Transformer model. We then replaced the original domain sequence in Q1H158 with the sampled sequences and compared the structures predicted with Alphafold based on the original and modified sequences. For comparison, we also sampled sequences from RITA using beam search. We note that one reason for choosing Q1H158 is that the domain we want to redesign is at the end of the sequence, enabling a causal language model like Rita to sample the domain conditioned on the rest of the protein. We show the results in Fig. 9, where several sequences sampled with Rita have a significantly lower TM-score than sequences sampled from the Transformer. A closer analysis showed that some of these sequences did not contain a domain recognized by the Pfam HMM for family PF00289, indicating the the fine-tuned RITA model did not always complete the sequence with the same domain as is found in the original sequence, as desired. While such alternative completions might very well correspond to a domain organization found in natural sequences, it shows that some care has to be taken when using unconditional language models for redesigning parts of a sequence, even if the model has been fine-tuned only with examples for the desired domain organization. On the other hand, the decoder of the shallow Transformer has been trained only for sampling the desired domain.

**Figure 9:**
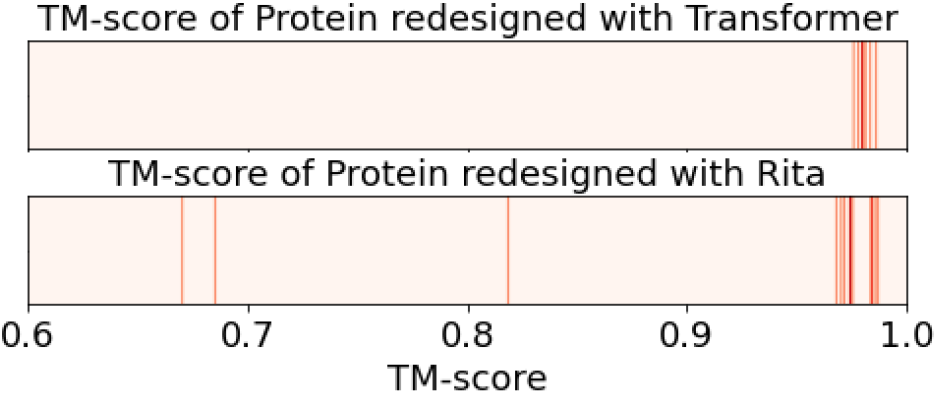
TM-scores comparing Alphafold structural predictions based on original and modified sequences of the protein Q1H158, which contains domains PF00289 and PF02785. The sequences are modified by resampling PF00289 from the shallow Transformer and RITA.

Finally we looked at a method for unsupervised structural prediction called Direct Coupling Analysis (DCA). We sampled for each input protein sequence of the training set 8 target sequences, adding all sampled sequences together with the input sequences to a pMSA, which therefore contained natural sequences from the input domain concatenated to artificial sequences from the target domain. We then attempted to extract structural contacts between the two domains using plmDCA [9], a popular method for prediction contacts. While the performance in contact prediction is worse than when using the natural target sequences directly, see Appendix Sec. D.2, there is a strong signal with several correctly predicted contacts among the highest-scoring residue pairs.

## 5 Discussion

In this work, we explored the use of Transformers for generating protein domain sequences while taking into account other domain sequences that are part of the same multi-domain protein. We cast the problem as a translation task, which allowed us to directly use Transformers developed for translation between natural languages. We showed that this architecture is capable of outperforming state-of-the-art shallow autoregressive models in several metrics and explored a new regularization scheme optimized for our use case. Casting the task as a translation problem allowed us to use metrics like the matching performance for assessing the quality of the generative models.

Our work is placed at the intersection of two streams of research: There is a long history of building domain specific generative models on aligned sequences for tasks like drug design or mutational effect prediction. More recently, however, large models based on Transformer architectures trained on all or nearly all unaligned protein sequences available have shown remarkable capabilities for capturing complex patterns in the data. Our work, on the other hand, solves a very generic sequence-to-sequence prediction task using smaller Transformer architectures, specialized for a family pair and using aligned sequences, which allows for domain-specific models. One limitation of our work is that we consider only a single domain as the context when predicting the sequence of an interacting domain, disregarding additional domains that might be present in the same protein. Conceptually, however, it would not be hard to create a model that takes more than one other domain as context when generating a new sequence.

An interesting question for further research is if we could solve the task equally well or even better using a single large model for training on all families at once. We note however, that this is not straightforward since a family might appear with more than one different family on the same protein, making the prediction task ambiguous. From the practical perspective, our method can be used for all tasks where a novel amino acid sequence needs to be generated conditioned on other amino acid sequences. This includes the case of general protein-protein interaction, which we did not analyze in the current work, but which can in principle be solved using the same models we presented.

## Appendix

### Appendix A Datasets

In this section we give details about the 27 family pairs used to measure the performance of the different models. The quantities *N*_*in*_ and *N*_*out*_ are the domain length of the source family and the target family, *M*_*train*_ and *M*_*val*_ the size of the training set and validation set and *d*_*med*_ is the median distance of a sequence in the validation set to the training set. This distance was used as a cutoff for distinguishing the matching performance for sequences close or far from the training set, which are denoted by ℳ_*Close*_ and ℳ_*Far*_.

### Appendix B Methods and Models

#### B.1 Transformer

The translation model we use is matching closely the original transformer model from Ref. [33], featuring an encoder-decoder architecture. While we refer to this work for more details, we review here the key components. Sequences from the source family are encoded by the encoder and used as the input for the decoder. The source sequence is processed through alternating blocks of self-attention and linear layers. The same is done for the already translated part of the target sequence, while the part of the target sequences not yet decoded is masked. Typical vocabulary sizes in NLP are in the order of 10^4^ to 10^5^, while in our case we have a vocabulary 𝒱 is composed of 21 tokens, corresponding to 20 amino acids and an alignment gap symbol.

The input embedding is composed of two parts, one for the amino acid identity and one for the position in the sequence. We learn a dictionary *W*, mapping each of the 21 symbols to a vector of dimension *d*_*model*_. The sequence position is embedded as a vector *PE*, calculated as

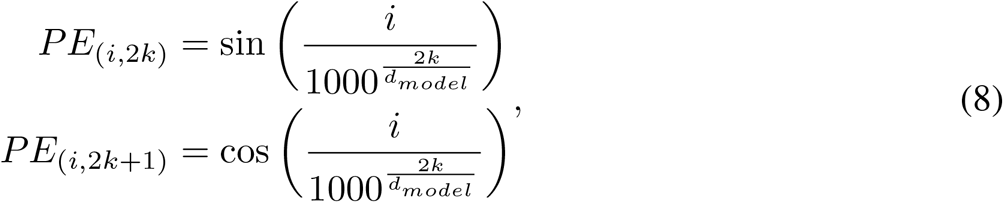

where *i* is the position in the sequence and *k* is dimension in the embedding vector. The embedding of a sequence is then taken as the sum of the amino acid and positional embeddings. The embedded amino acid sequences are then passed to the encoder, mapping them to a latent representation *z* = (*z*_1_, …, *z*_*n*_). This latent representation is then passed to the decoder that predicts the interaction partner sequence.

**Table 1:** List of the pairs of domains used in this dataset. *N*_*in*_ and *N*_*out*_ are the length of the input domain and the target domain. *M*_*train*_ and *M*_*val*_ and the size of the training and validation dataset.

| Source Family | Target Family | $N_{in}$ | $N_{out}$ | $M_{train}$ | $M_{val}$ | $d_{med}$ |
| --- | --- | --- | --- | --- | --- | --- |
| PF00689 | PF00122 | 184 | 223 | 4460 | 488 | 62 |
| PF00289 | PF02786 | 112 | 213 | 4502 | 495 | 35 |
| PF02785 | PF00289 | 110 | 112 | 4396 | 490 | 24 |
| PF00004 | PF07724 | 134 | 173 | 4886 | 1118 | 21 |
| PF00006 | PF02874 | 215 | 71 | 5370 | 464 | 15 |
| PF02785 | PF02786 | 110 | 213 | 4542 | 494 | 35 |
| PF00207 | PF07677 | 94 | 94 | 552 | 138 | 10 |
| PF00207 | PF01835 | 94 | 98 | 1027 | 241 | 43 |
| PF08264 | PF00133 | 154 | 604 | 5846 | 501 | 110 |
| PF00501 | PF13193 | 425 | 76 | 17050 | 498 | 120 |
| PF01591 | PF00300 | 225 | 196 | 760 | 189 | 31 |
| PF08240 | PF00107 | 111 | 132 | 16903 | 500 | 38 |
| PF02770 | PF02771 | 99 | 115 | 12048 | 499 | 33 |
| PF00441 | PF02771 | 152 | 115 | 12703 | 501 | 38 |
| PF01842 | PF13840 | 69 | 67 | 851 | 190 | 19 |
| PF08545 | PF08541 | 82 | 92 | 2548 | 556 | 35 |
| PF00441 | PF02770 | 152 | 99 | 12850 | 497 | 26 |
| PF00005 | PF08402 | 139 | 78 | 4933 | 1066 | 49 |
| PF00005 | PF08352 | 139 | 67 | 5751 | 1243 | 40 |
| PF00664 | PF00005 | 276 | 139 | 15889 | 3473 | 106 |
| PF03171 | PF14226 | 103 | 120 | 5624 | 1317 | 29 |
| PF12780 | PF12781 | 270 | 222 | 1498 | 361 | 37 |
| PF12775 | PF12780 | 274 | 270 | 1632 | 386 | 39 |
| PF07724 | PF10431 | 173 | 83 | 7315 | 1649 | 25 |
| PF00690 | PF00702 | 71 | 212 | 4450 | 486 | 63 |
| PF00690 | PF00122 | 71 | 223 | 5356 | 484 | 37 |
| PF00004 | PF10431 | 134 | 83 | 4685 | 1088 | 25 |

The decoder implements an auto-regressive distribution

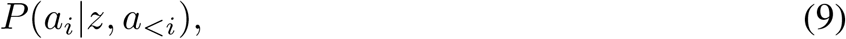

defining the probability of the *i*^*th*^ amino acid in the interaction partner sequence given the preceding amino acids in the interaction partner *a*_*<i*_ and the hidden representation *z* of the input sequence *B*. During training we use the true amino acids for *a*_*<i*_, while during sampling we sample the sequence *A* sequentially.

##### Attention Mechanism

Both the encoder and decoder use self-attention mechanisms and the decoder also the cross-attention mechanism.

Following [33], we define the attention operation as

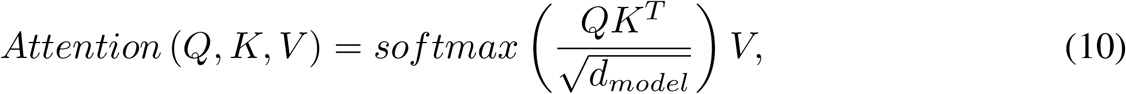

where *Q* is the query matrix, *K* is the key matrix, *V* is the value matrix and *d*_*model*_ the dimension of the keys, which we will define below.

For each element of the query *Q* we compute its similarity with the different values of the keys *K*. This yields weights used to compute a weighted average of the value *V*.

The output of the *i*^*th*^ attention head, called *head*_*i*_, is calculated as

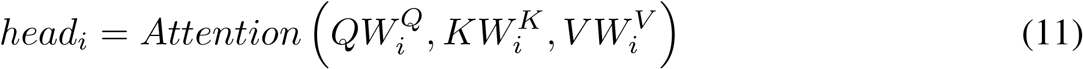

We then linearly combine the different heads, resulting in

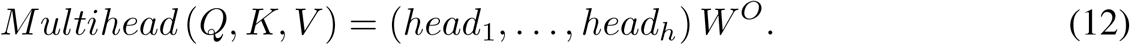

In self attention layers, the keys, queries and values are calculated from the same input. In this case, *K* = *Q* = *V*.

In cross attention, the keys and values are based on *z* and the queries on the intermediate decoder representations. In the results presented in this paper we only used models with a single head.

##### Transformer Architecture

The complete architecture is represented in Fig. B.1 and is based on stacking encoder and decoder blocks in the two parts. At the end of these blocks, linear with dimension *d*_*ff*_ and residual connections to the input of the blocks are added.

**Figure B.1:**
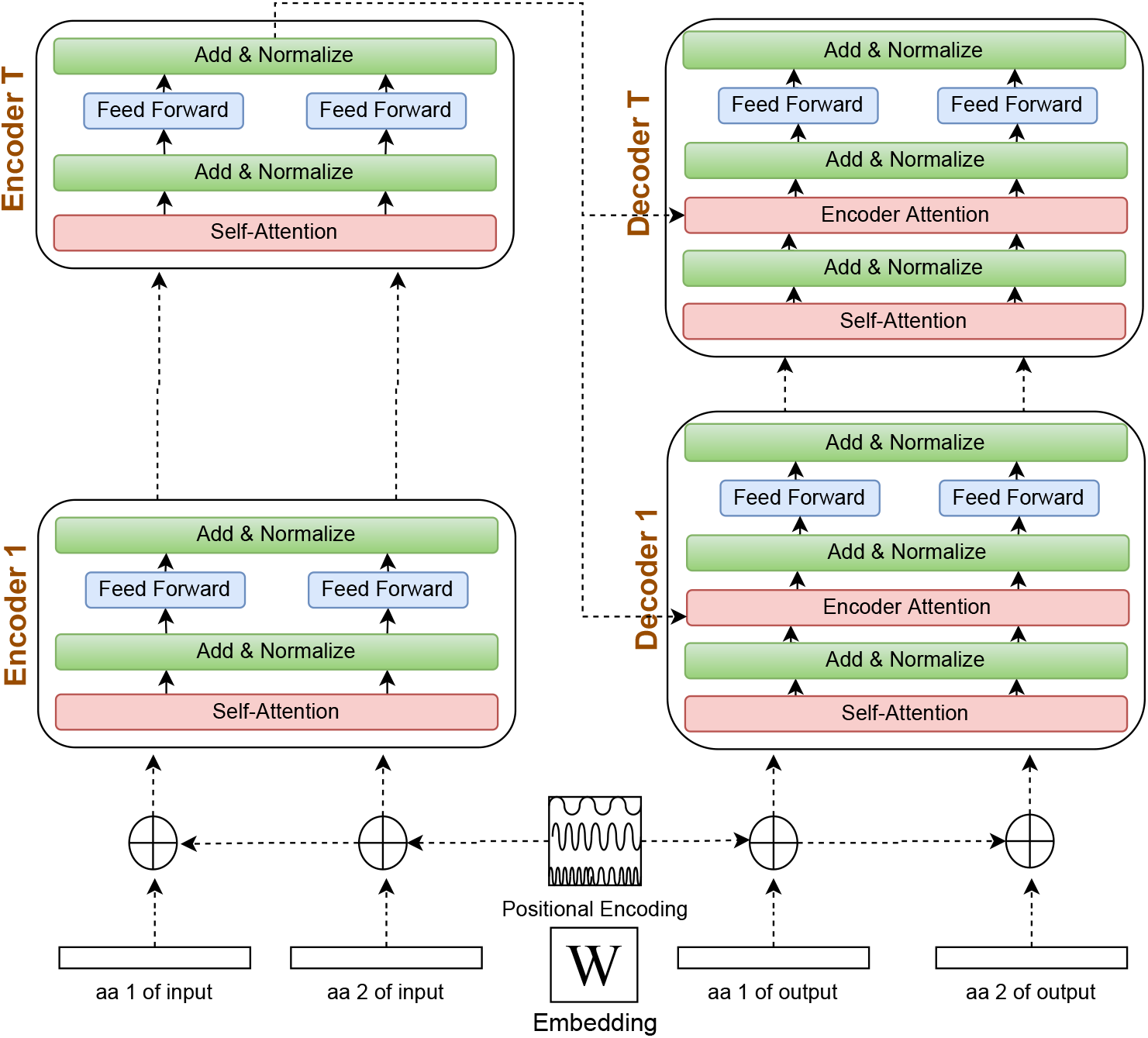
Transformer architecture for an input and output protein of length 2 and T layers: *W* is an embedding matrix, matching each amino acid to a vector of size *e*. The architecture here represents proteins of length 2 for simplicity, but the encoder and decoder can handle inputs of arbitrary length. The positional encoding uses sine functions of different frequencies to generate an embedding of the protein position.

The code is based on the the PyTorch implementation of the Transformer https://github.com/pytorch/pytorch.

#### B.2 arDCA Baseline

As a baseline we use the recently introduced arDCA [32], which is an efficient autoregressive model for protein sequences. or an amino acid sequence *A* = (*a*_*i*_, …, *a*_*N*_) of length *N*, arDCA defines the conditional probability *P* (*a*_*i*_|*a*_*i−*1_, …, *a*_1_) as

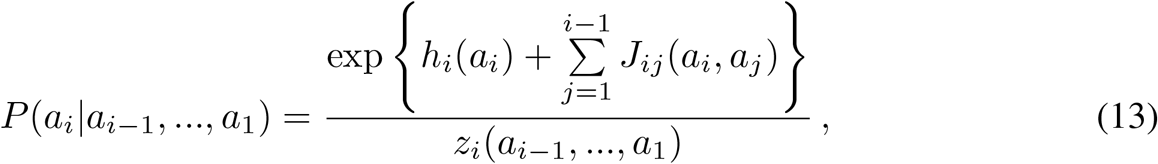

with 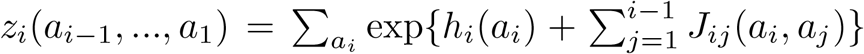 being a normalization factor. The parameters *h* depend on a single position and the amino acid found at that position and the parameters *J* on pairs of positions and the two amino acids found at that pair of positions.

The probability of a sequence can be computed using the decomposition

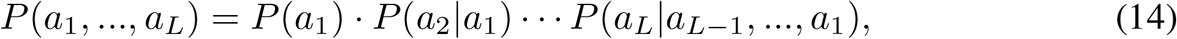

which is tractable. Training can be done using standard convex optimization methods.

For our purposes, we concatenate the source protein *B* and the target protein *A* into a single sequence during training. During evaluation, we just need the conditional probability *P* (*A*|*B*), which we calculate using

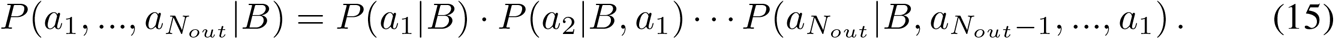

We also added an L2 regularization on the parameters *h* and *J*. During our experiments we used the regularization parameters communicated by the authors (*λ*_*h*_ = *λ*_*J*_ = 0.0001).

#### B.3 RITA

Rita is decoder only Transformer without conditioning information. It means that is only use the decoder part of B.1, where encoder-decoder attention layer has been deleted. This defines a generic autoregressive models. We used the Rita L model composed of Large 680*M* parameters, a model dimension of 1536, and 24 layers. This huge model was then trained on the UniRef-100 database. Moreover we should note that the training was done in both direction. This explains why we presents both score when evaluating the performance of Rita finetuned. For finetuning Rita we used a batch-size of 6 and the Adam optimizer on the full length sequence (both direction) of the proteins in our train-set. Every Validation loss was computed every 200 gradient updates, and the best performing model was kept for the experiments shown in the paper.

Rita is a language model based on the decoder of the original Transformer model in Fig.B.1. This means it does not use encoder-decoder attention and implements a generic unconditioned autoregressive sequence model. In our experiments, we use Rita L, which has 680*M* parameters, a model dimension of 1536 and 24 layers. The model we used for finetuning was pre-trained on Uniref-100 predicting both in the natural and the reverse direction of the protein sequences. We used a batch-size of 6 and the Adam optimizer for finetuning on the full length sequences (both directions) on our datasets. We calculated the loss on the validation every 200 gradient updates on a total of 4000 gradient updates. We used the best performing model for the experiments shown in the paper. The number of steps to finetuned each model vary between families, but usualy stand around 1000, way before the end of our training. To make the loss comparable we only took into consideration the position that were match state for the Pfam HMM of the domain.

#### B.4 Training Time

In this section we present a table with the training time of our models using a single Nvidia V100 GPU. The first column refers to the family pair, the second to the training time of the shallow Transformer, the third one to the large Transformer with entropic regularization and the last one to the large Transformer without entropic regularization.

**Figure C.1:**
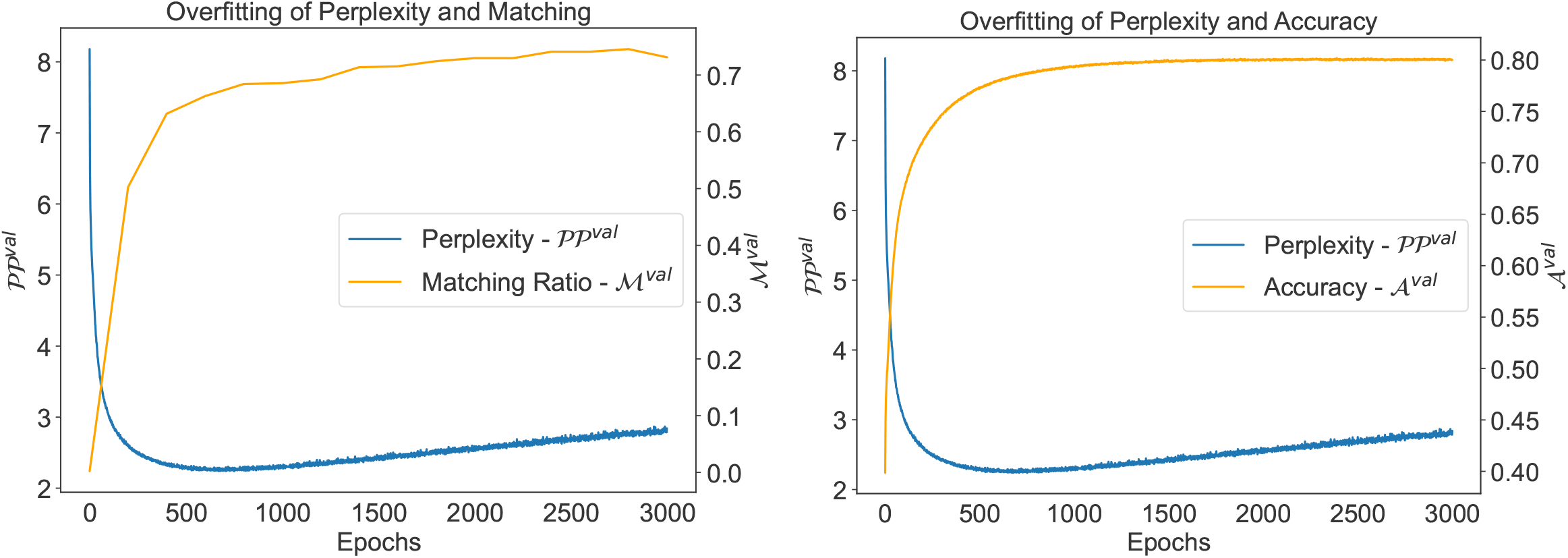
Evolution of the perplexity and the number of correctly matched pairs (a) on the validation set and (b) the accuracy during training for PF013171-PF14226 for the large transformer.

| Pair | Runtime Shallow | Runtime Renyi | Runtime Large |
| --- | --- | --- | --- |
| PF00289_PF02786 | 0 days 10:13:10 | 3 days 17:10:52 | 0 days 15:10:43 |
| PF02785_PF02786 | 0 days 10:20:21 | 2 days 10:45:01 | 0 days 11:24:58 |
| PF02785_PF00289 | 0 days 09:50:34 | 2 days 02:05:06 | 0 days 15:45:50 |
| PF00004_PF07724 | 0 days 19:54:55 | 2 days 15:57:20 | 0 days 23:05:37 |
| PF00006_PF02874 | 0 days 12:00:52 | 2 days 07:07:35 | 0 days 12:28:15 |
| PF00207_PF07677 | 0 days 01:44:11 | 0 days 06:14:45 | 0 days 01:09:31 |
| PF00207_PF01835 | 0 days 03:30:15 | 0 days 11:15:13 | 0 days 02:17:15 |
| PF08264_PF00133 | 0 days 18:21:48 | 6 days 05:37:06 | 0 days 19:24:28 |
| PF00501_PF13193 | 1 days 16:47:12 | 12 days 12:27:26 | 1 days 07:23:54 |
| PF01591_PF00300 | 0 days 03:20:45 | 0 days 19:01:09 | 0 days 02:23:11 |
| PF08240_PF00107 | 1 days 13:11:32 | 6 days 20:56:44 | 1 days 18:54:09 |
| PF02770_PF02771 | 1 days 02:09:05 | 6 days 01:37:54 | 1 days 07:02:04 |
| PF00441_PF02771 | 1 days 01:16:55 | 10 days 18:35:05 | 1 days 08:18:41 |
| PF00441_PF02770 | 1 days 02:20:40 | 6 days 01:56:31 | 1 days 11:05:05 |
| PF01842_PF13840 | 0 days 02:38:04 | 0 days 10:54:23 | 0 days 02:37:06 |
| PF08545_PF08541 | 0 days 07:48:42 | 1 days 04:28:05 | 0 days 05:54:03 |
| PF00005_PF08402 | 0 days 11:30:09 | 1 days 19:43:05 | 0 days 12:58:03 |
| PF00005_PF08352 | 0 days 13:42:05 | 2 days 13:55:30 | 0 days 16:48:21 |
| PF00664_PF00005 | 2 days 01:30:23 | 10 days 19:21:16 | 2 days 02:02:05 |
| PF03171_PF14226 | 0 days 22:10:30 | 4 days 04:10:56 | 0 days 21:30:18 |
| PF12780_PF12781 | 0 days 07:10:21 | 1 days 00:33:05 | 0 days 03:09:31 |
| PF12775_PF12780 | 0 days 08:13:18 | 2 days 05:13:49 | 0 days 03:43:15 |
| PF07724_PF10431 | 0 days 18:03:28 | 3 days 14:18:21 | 1 days 08:33:09 |
| PF00690_PF00702 | 0 days 09:55:26 | 2 days 16:45:33 | 0 days 10:26:16 |
| PF00690_PF00122 | 0 days 12:05:13 | 3 days 04:58:34 | 0 days 16:48:12 |
| PF00689_PF00122 | 0 days 10:37:22 | 5 days 08:58:05 | 0 days 17:20:30 |
| PF00004_PF10431 | 0 days 13:00:08 | 2 days 19:41:44 | 0 days 18:51:31 |

### Appendix C Regularization

#### C.1 Dropout and Weight-Decay Benchmark

In this section we show learning curves related to overfitting behaviour and regularization.

**Figure C.2:**
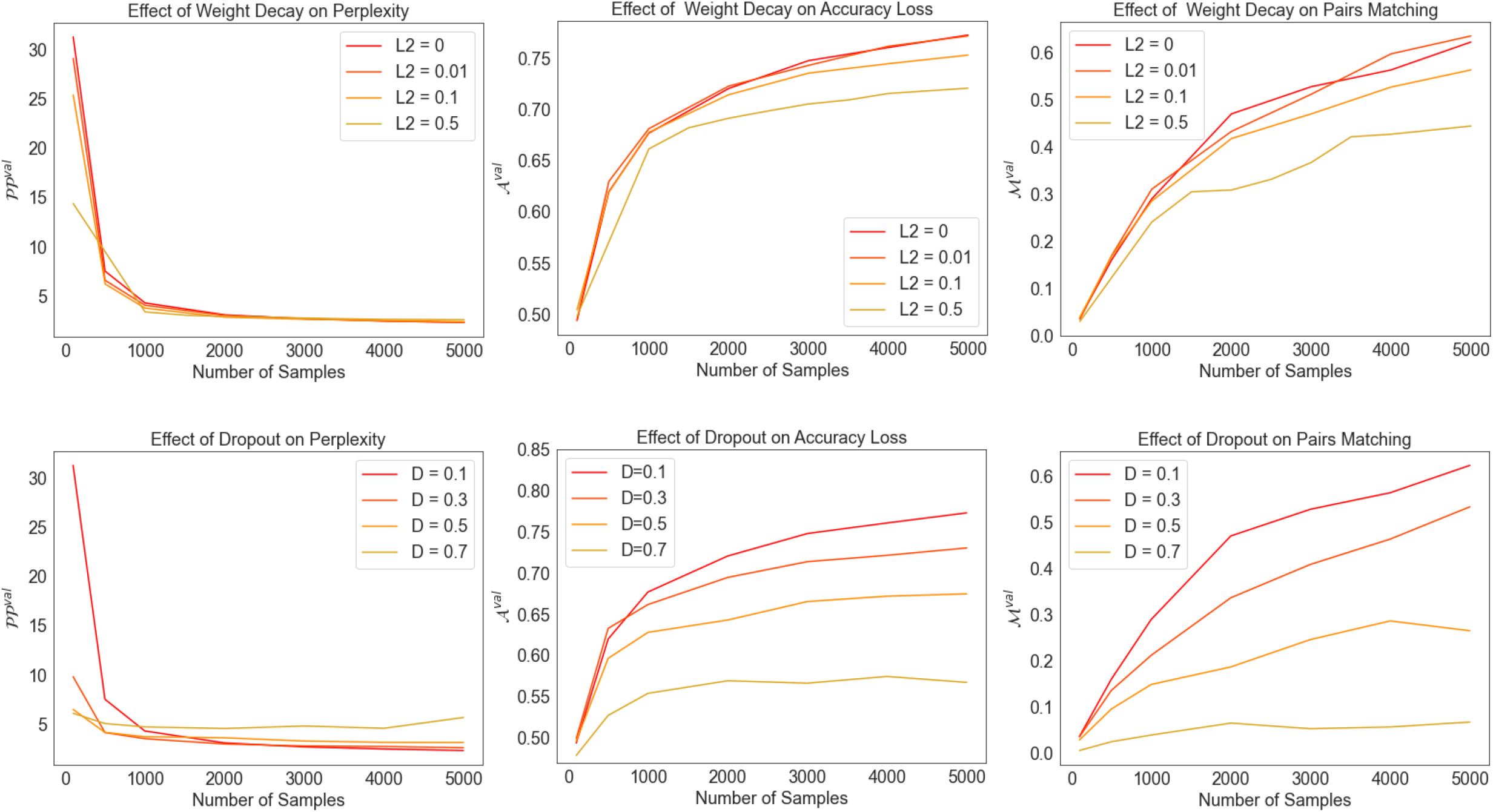
Performance of the Transformer on PF013171-PF14226 for varying number of training examples: Left plots show the perplexity 𝒫𝒫^*val*^, center plots show the accuracy 𝒜^*val*^, right plots show the ratio of correctly matched pairs ℳ^*val*^. All values are for the validation set.

#### C.2 Entropic Formulation of The Regularization

There is a relation between the entropic regularization and the Rényi entropy: For a number of sampled sequences *S* sufficiently large we can rewrite the regularization term

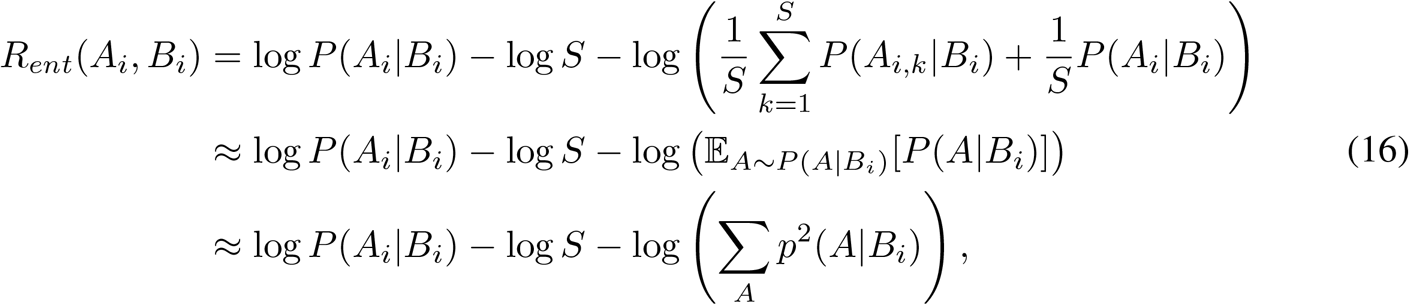

where the last sum is over all possible sequences *A*. The first term in the last line is equivalent to the standard loss and can be absorbed there. The second term is a constant and will not influence the gradient. The last term is the logarithm of the Rényi entropy of order 2, also called the *collision entropy*, of the distribution over target sequences conditioned on *B*_*i*_.

##### C.2.1 Entropic Regularization Performance

This section presents the results of the large Transformer trained with *α* = 0.7, and *S* = 5 in comparison with arDCA and the shallow Transformer in Fig. C.3, Fig. C.4 and Fig. C.5. The comparison with the large Transformer without regularization or with weight decay is presented in the following section, see Appendix Sec. C.3.

**Figure C.3:**
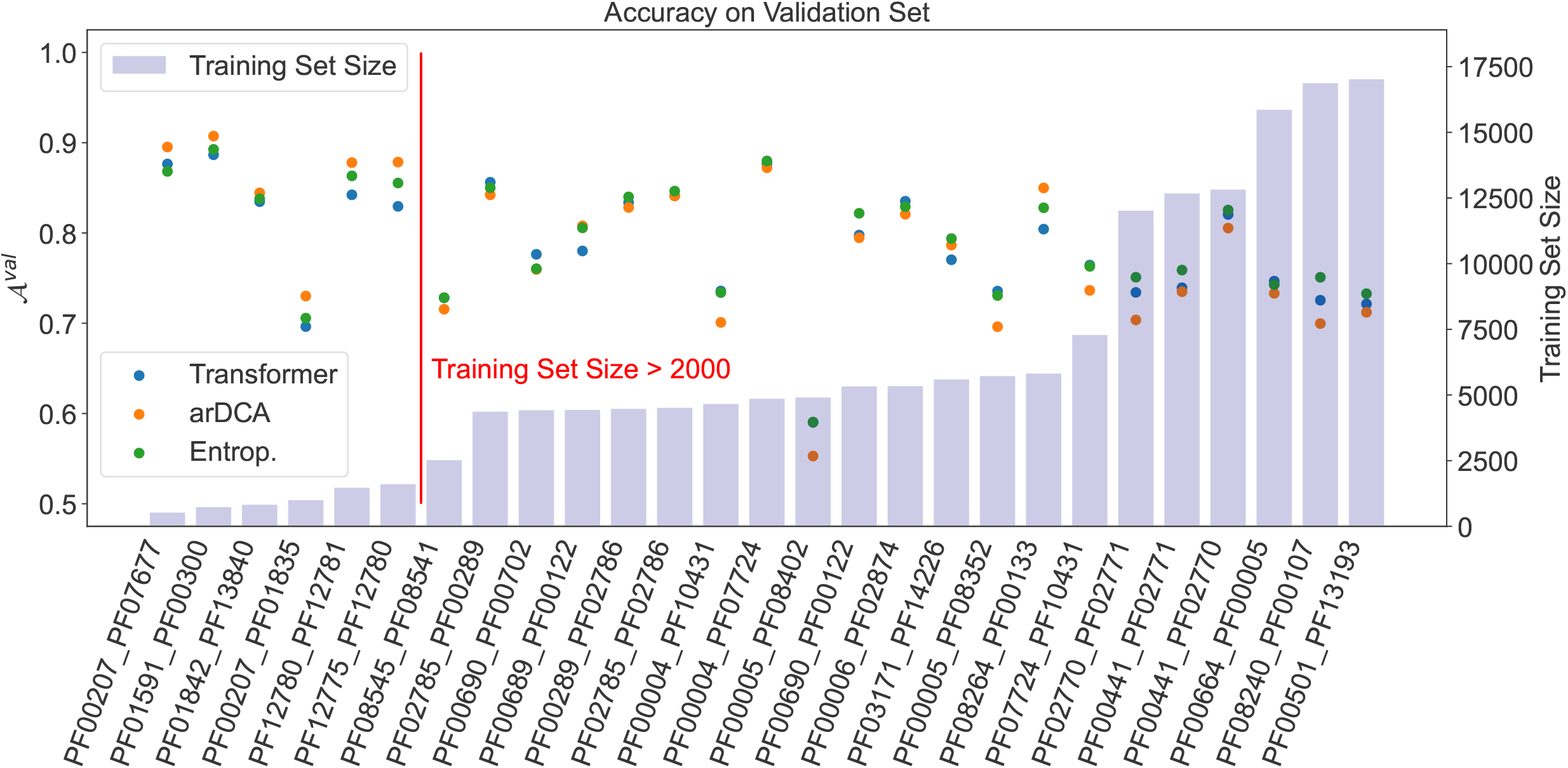
Accuracy 𝒜^*val*^ on the validation set for the shallow Transformer, arDCA, and the large Transformer with entropic regularization. The families are ordered by training set size.

**Figure C.4:**
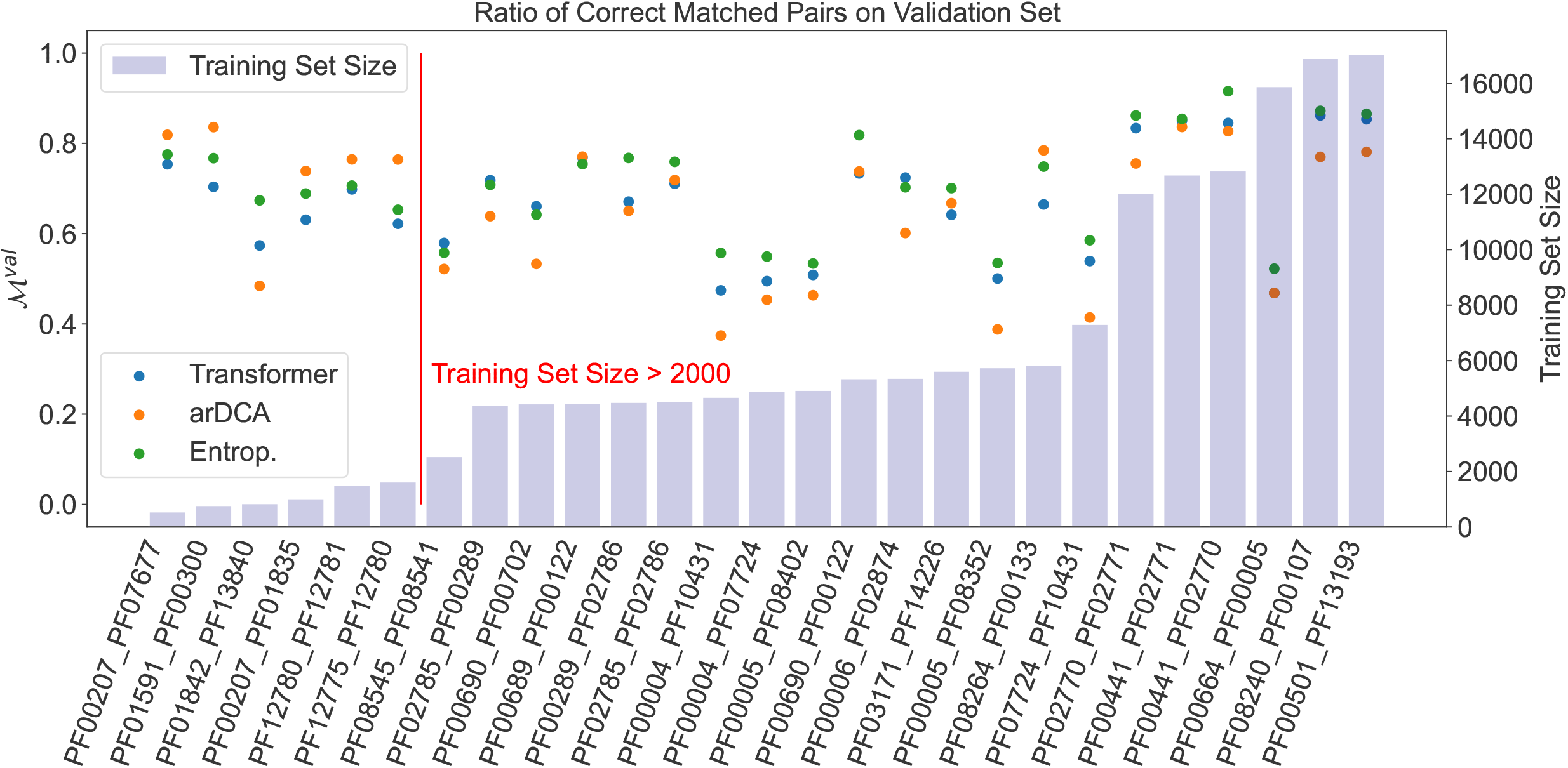
True positive rate for matching on the validation set for the shallow Transformer, arDCA, and the large Transformer with entropic regularization. The families are ordered by training set size.

**Figure C.5:**
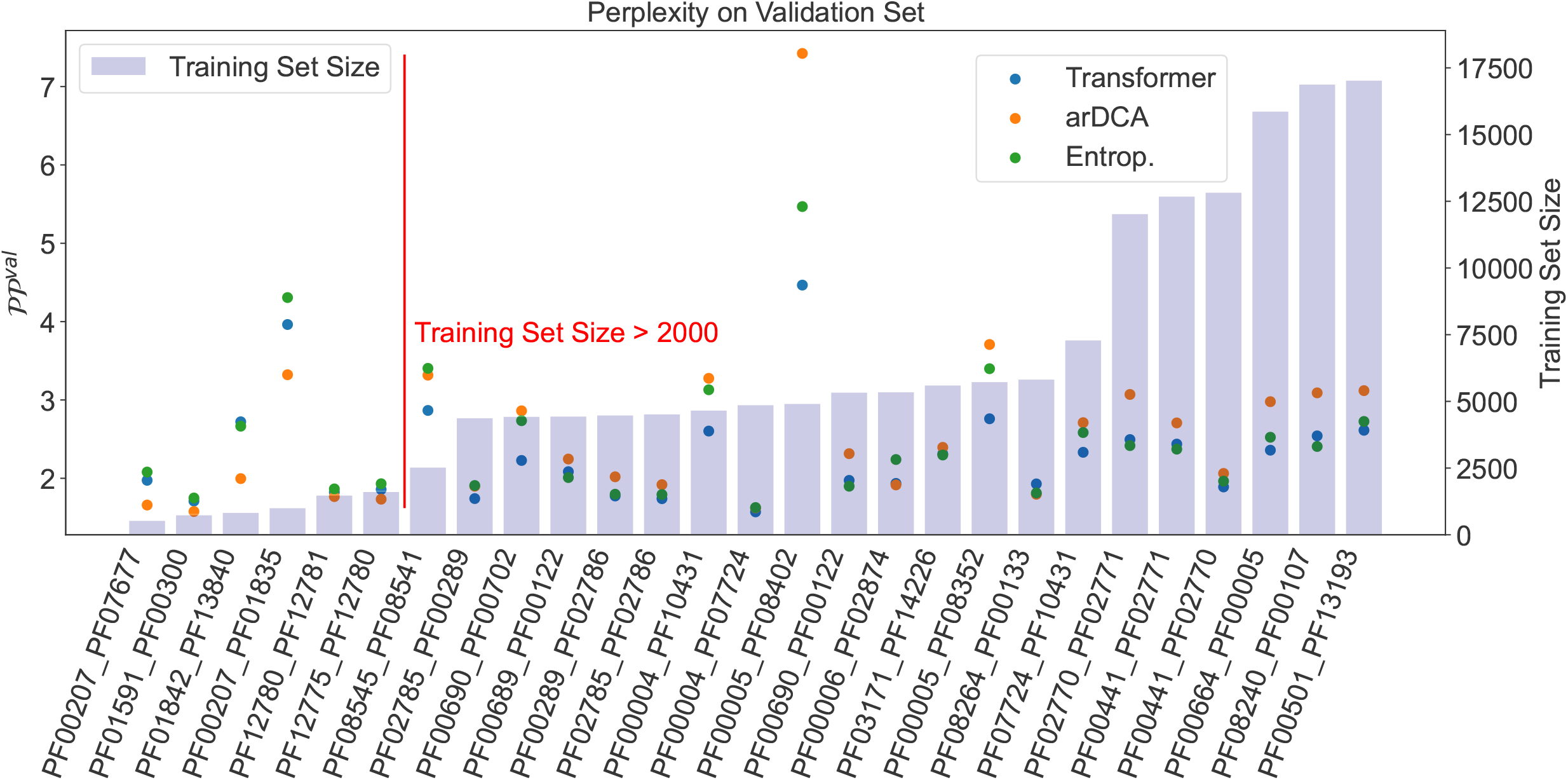
Perplexity 𝒫𝒫^*val*^ for the shallow Transformer, the large Transformer with entropic regularization and arDCA on the validation set. The families are ordered by training set size.

#### C.3 Entropic Regularization Compared with Weight Decay

In this section, we compare the entropic regularization and weight decay on different metrics, average over all 27 families.

**Figure C.6:**
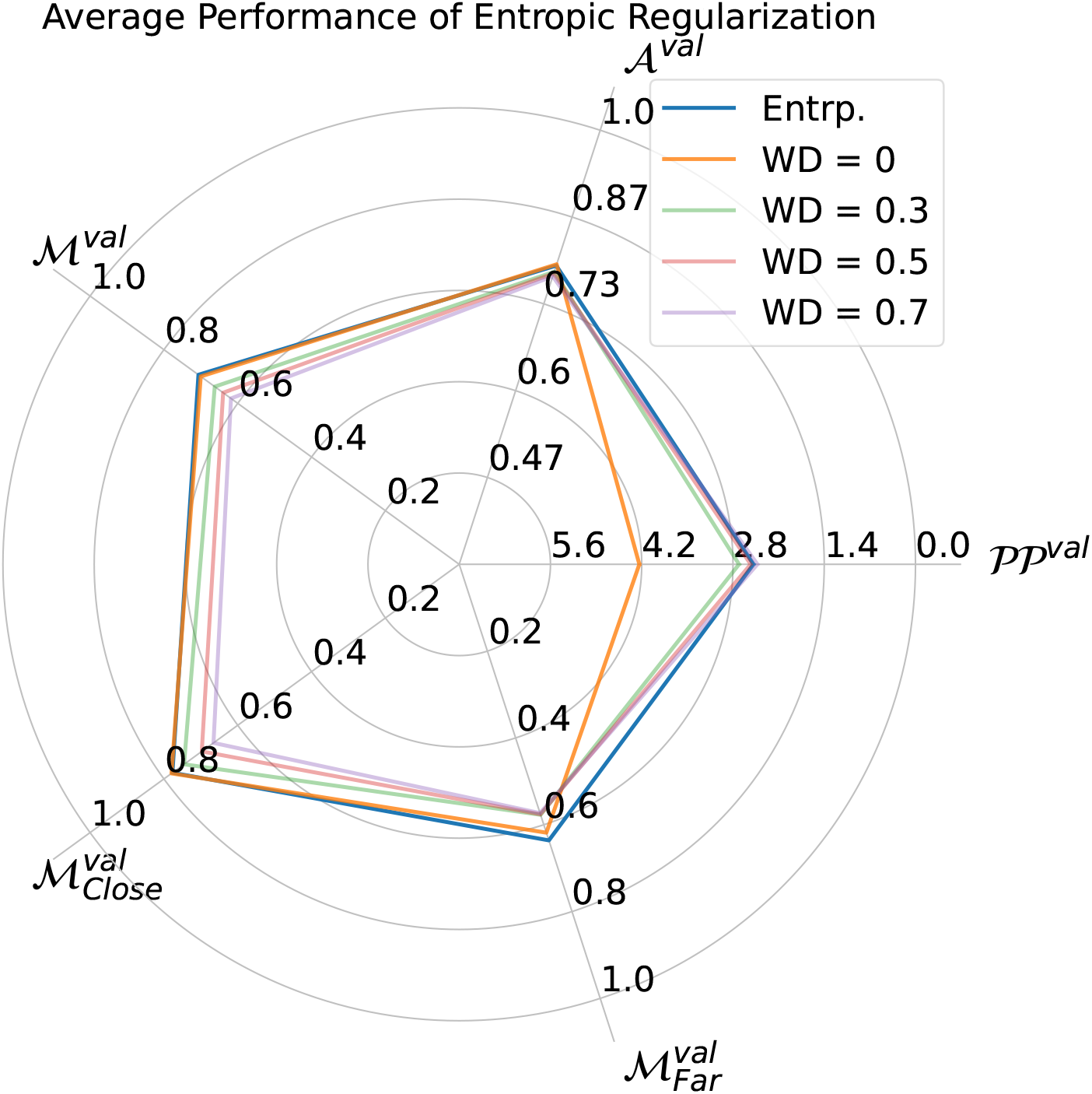
Radar plot comparing different regularization schemes, entropic (Entrp.) and weight decay (WD) with different strengths for the large Transformer. The radial direction of the perplexity 𝒫𝒫 is reversed in order to have the same direction for increasing performance as for the other metrics. The plot was done by averaging the metrics of all families.

##### Entropic versus Weight Decay

In this section we compare the performance of the large Transformer on different metrics for all families individually. The resulting plots were split into different figures in order to make them fit on the pages.

**Figure C.7:**
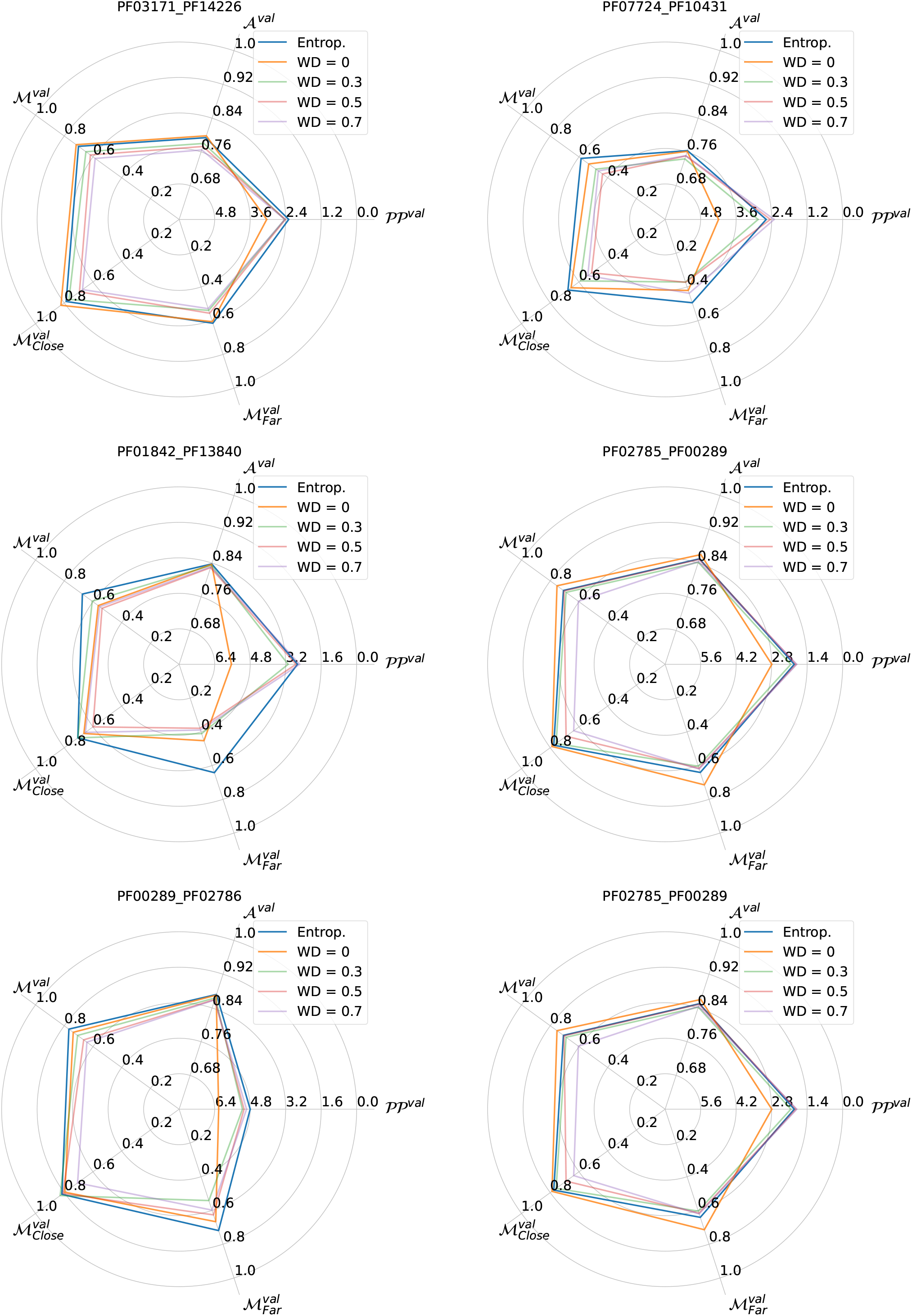
Comparing the performance of the large Transformer with entropic regularization and weight decay (WD). The radial direction of the perplexity 𝒫𝒫 is reversed in order to have the same direction for increasing performance as for the other metrics.

**Figure C.8:**
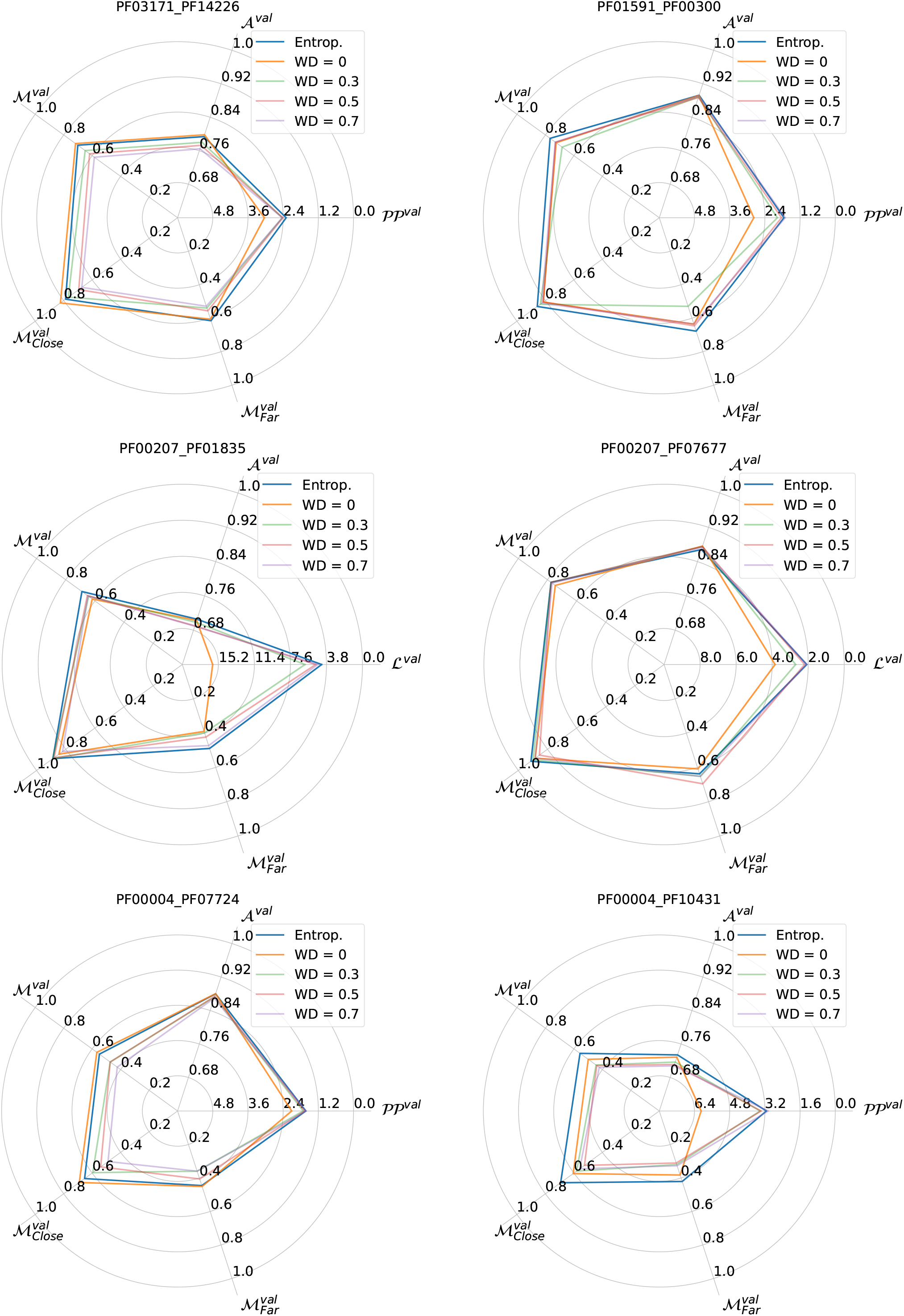
Comparing the performance of the large Transformer with entropic regularization and weight decay (WD). The radial direction of the perplexity 𝒫𝒫 is reversed in order to have the same direction for increasing performance as for the other metrics.

**Figure C.9:**
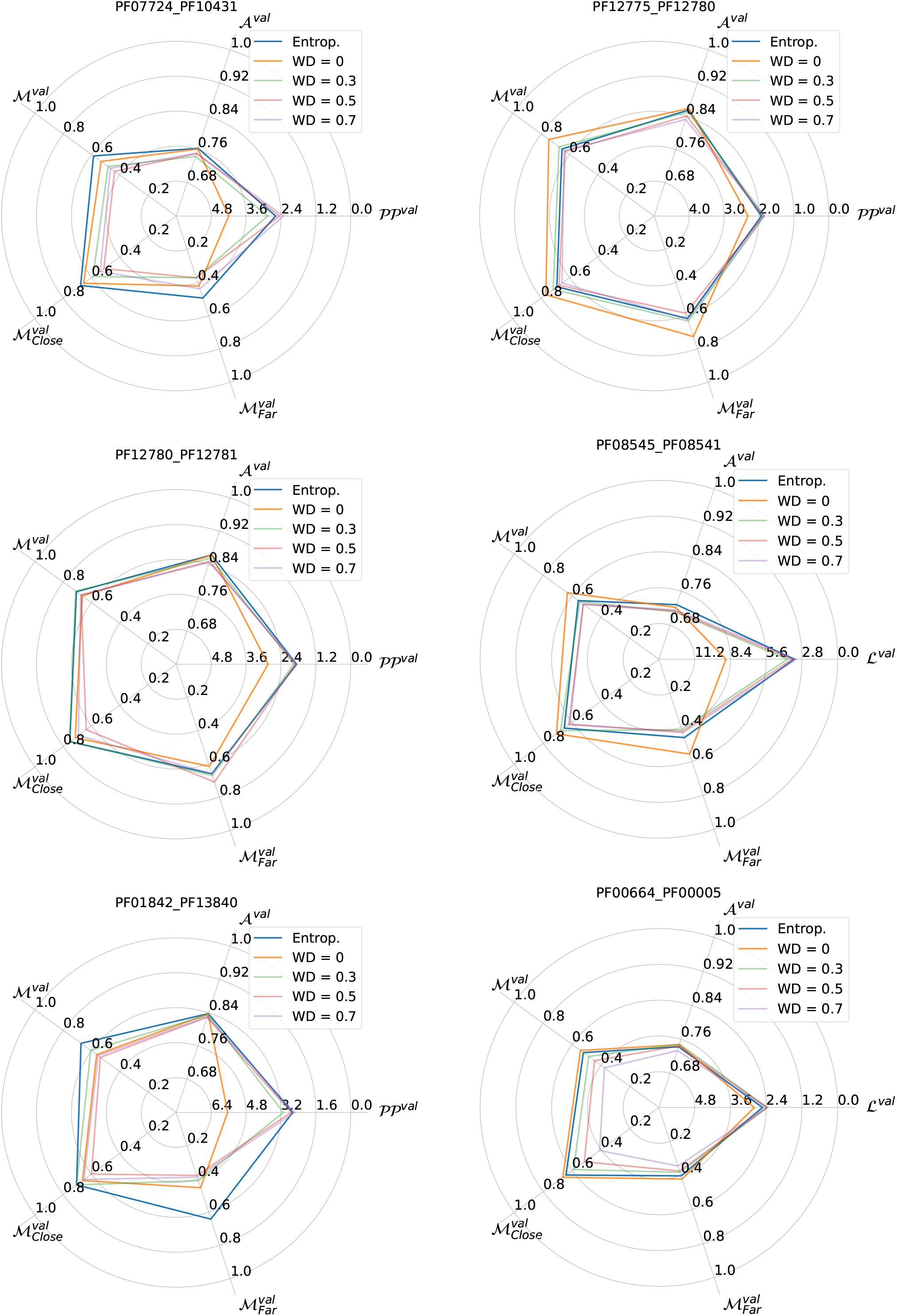
Comparing the performance of the large Transformer with entropic regularization and weight decay (WD). The radial direction of the perplexity 𝒫𝒫 is reversed in order to have the same direction for increasing performance as for the other metrics.

**Figure C.10:**
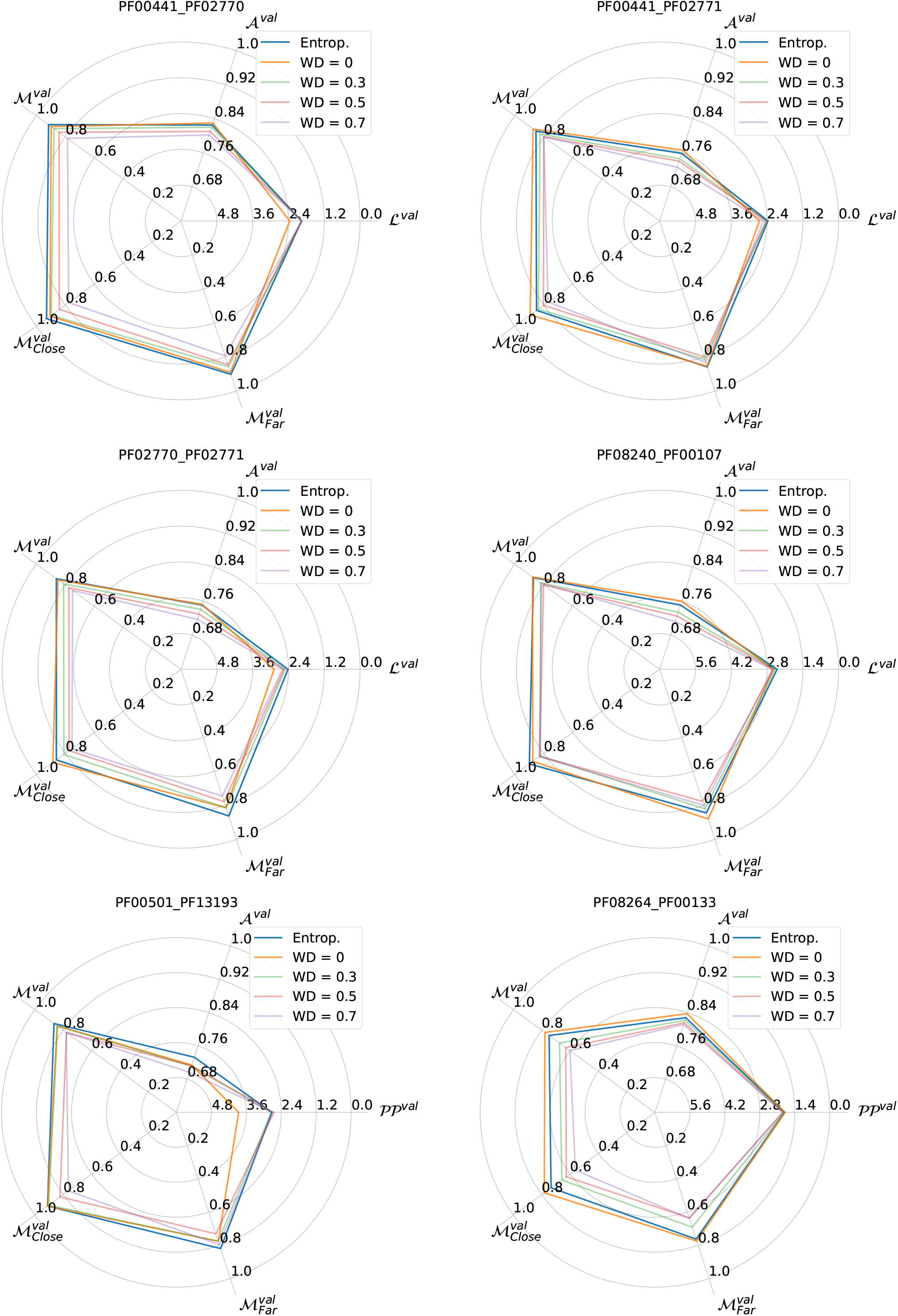
Comparing the performance of the large Transformer with entropic regularization and weight decay (WD). The radial direction of the perplexity 𝒫𝒫 is reversed in order to have the same direction for increasing performance as for the other metrics.

**Figure C.11:**
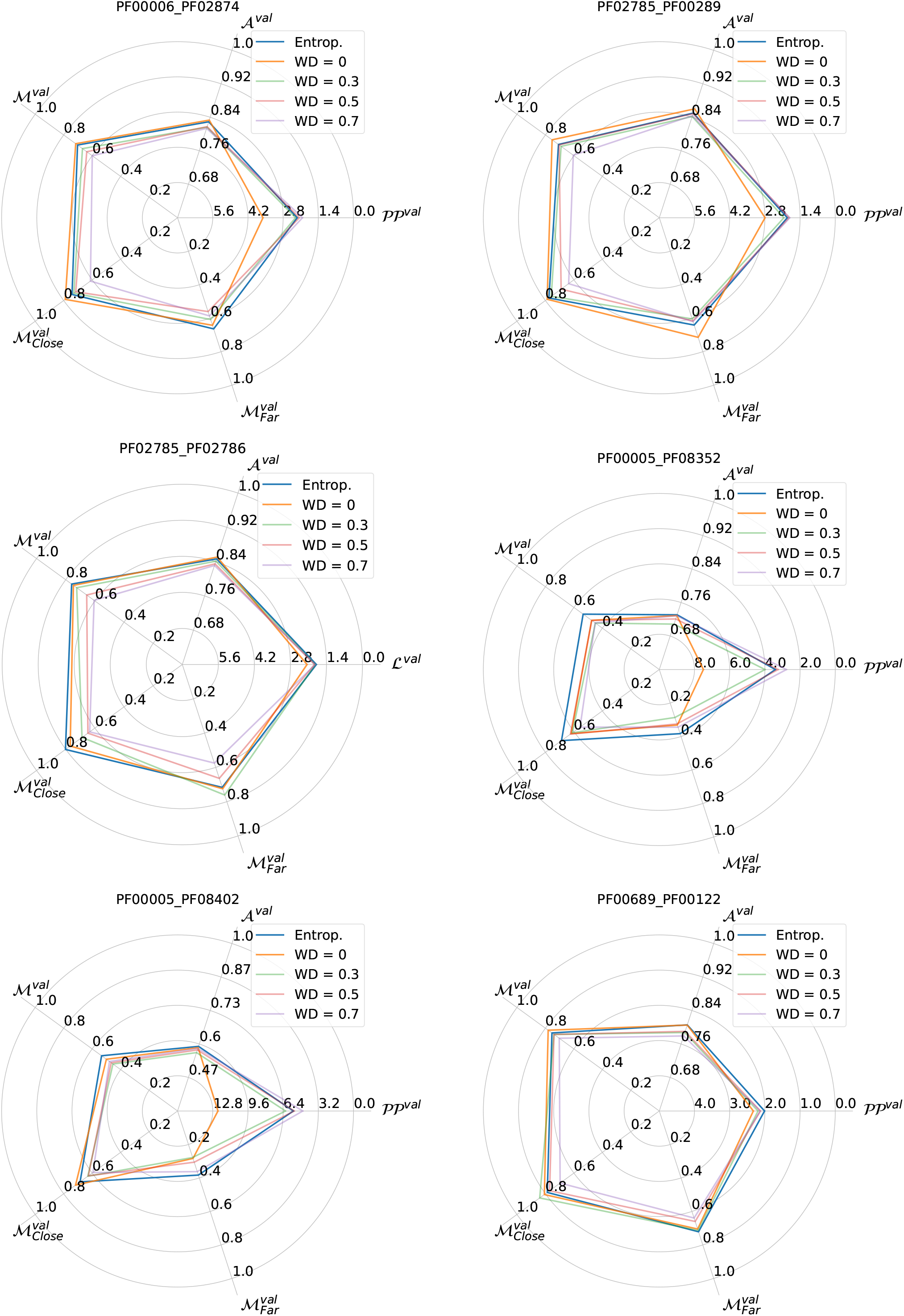
Comparing the performance of the large Transformer with entropic regularization and weight decay (WD). The radial direction of the perplexity 𝒫𝒫 is reversed in order to have the same direction for increasing performance as for the other metrics.

**Figure C.12:**
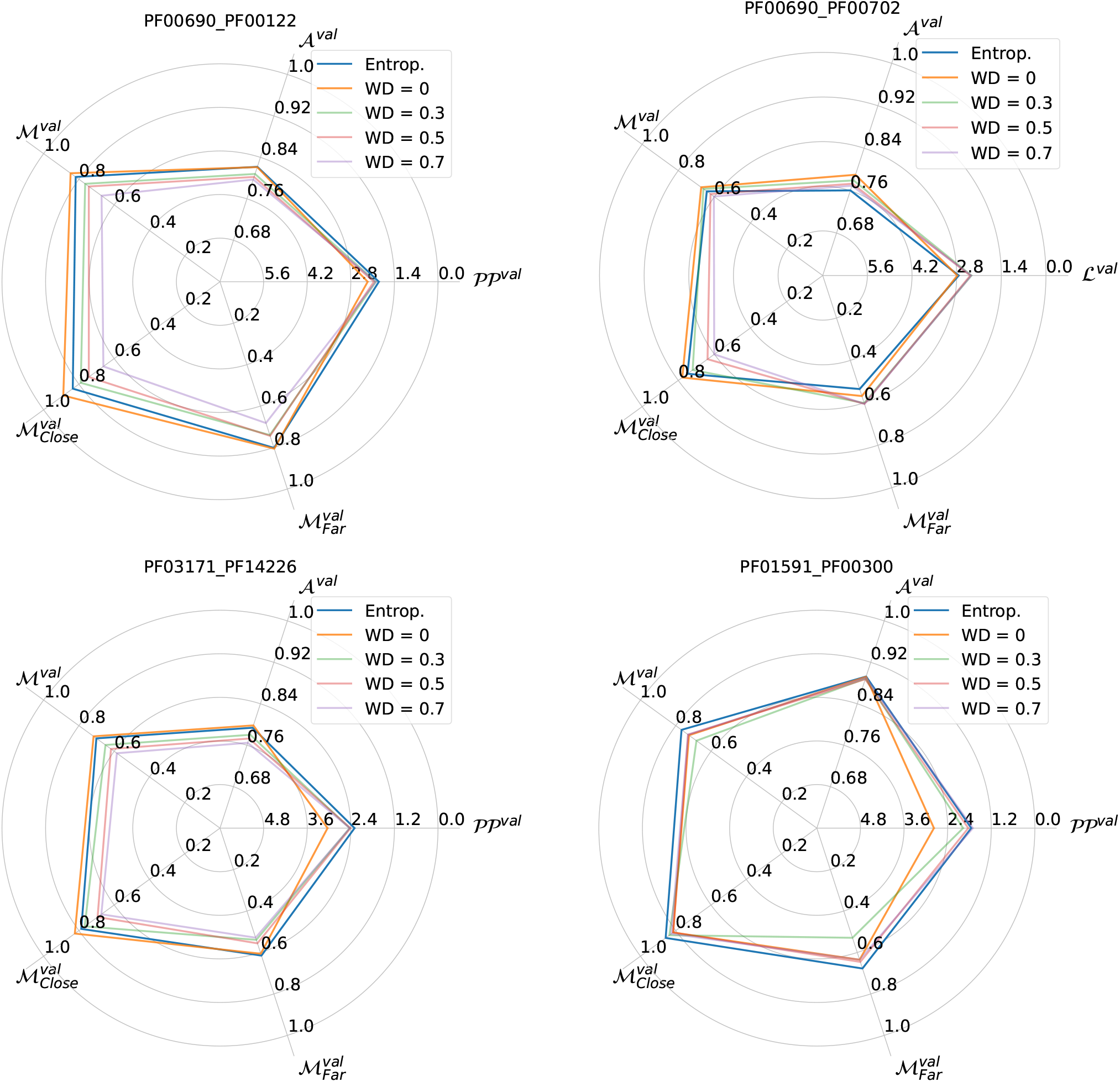
Comparing the performance of the large Transformer with entropic regularization and weight decay (WD). The radial direction of the perplexity 𝒫𝒫 is reversed in order to have the same direction for increasing performance as for the other metrics.

### Appendix D Additional Structural Results

#### D.1 Structural comparison using AlphaFold

For all our computation we used the implementation of CollabFold with template search, 5 models, and 3 recycle. [23].

**Figure D.1:**
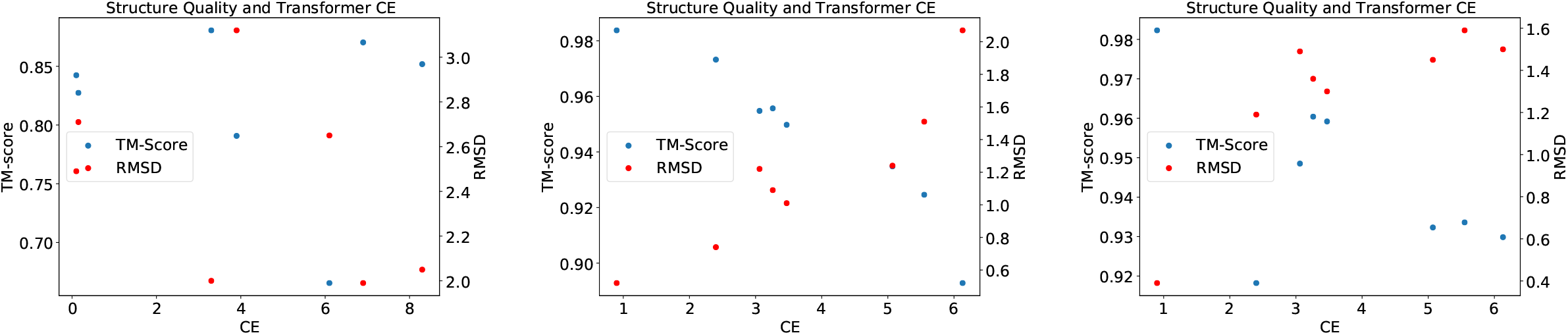
TM-Scores and RMSD values when comparing the Alphafold predicted structures of true sequences with Alphafold predicted structures of sequences where single domains have been replaced with homologous natural sequences. (Left is based on G0S4G4, which contains domains PF00004 and PF07724, which are in contact in PDB 5D4W. Homologous sequences are sampled from the validation set and inserted into the G0S4G4 sequence. Center is based on Q8ZN46, which contains domains PF00207 and PF01835, which are in contact in PDB 4U4J. Homologous sequences are sampled from the validation set and inserted into the Q8ZN46 sequence. Right is based on Q13SV4, which contains domains PF08545 and PF08541, which are in contact in PDB 4EFI. Homologous sequences are sampled from the validation set and inserted into the Q13SV4 sequence. In all of this proteins we measure the change in structural scores and in cross-entropy in the shallow Transformer model (abscissa).

#### D.2 Structural Information using DCA

Direct Coupling Analysis is a group of unsupervised methods for modelling aligned protein sequences, see Ref. [7]. Apart from other applications, it can be used for predicting structural contacts from MSAs.

##### plmDCA

plmDCA is a specific method of DCA based on a pseudolikelihood approximation for training the Potts Model. In this paper we used the asymmetric version of the method from https://github.com/pagnani/ArDCA.jl with default hyperparamters. The sequences sampled from the Transformer where realigned using HMMer [11].

##### Results per Pairs

In this section, we show the contact prediction results obtained with plmDCA for the 27 families.

**Figure D.2:**
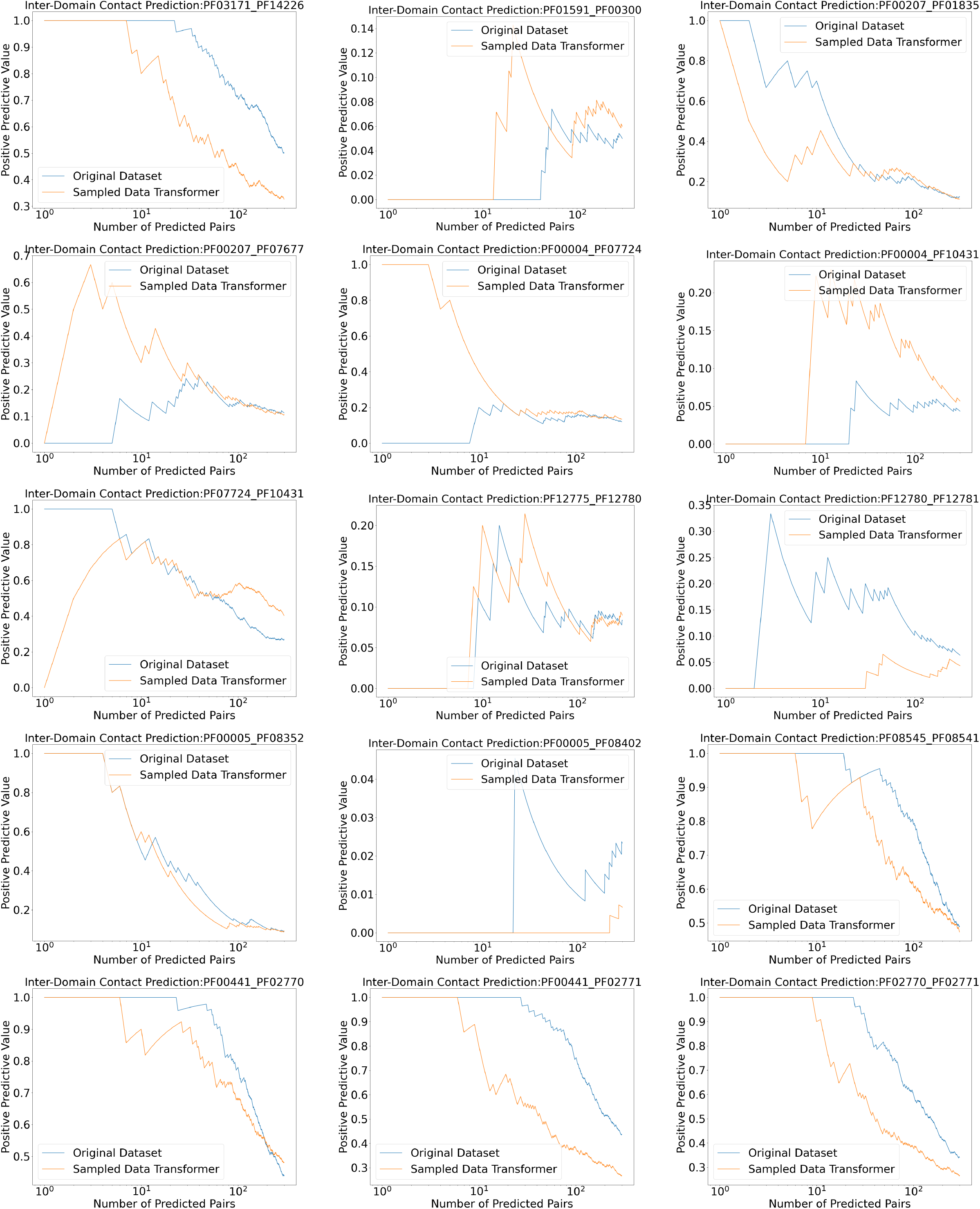
Contact Prediction using plmDCA for the original training set, and a sampled dataset from the the shallow Transformer. The curves represent the Positive Predictive Value (fraction of true positives) with respect to the number of predicted contacts. To make it fit the page format, we splitted the results on the different families in two figures: this one and the following.

**Figure D.3:**
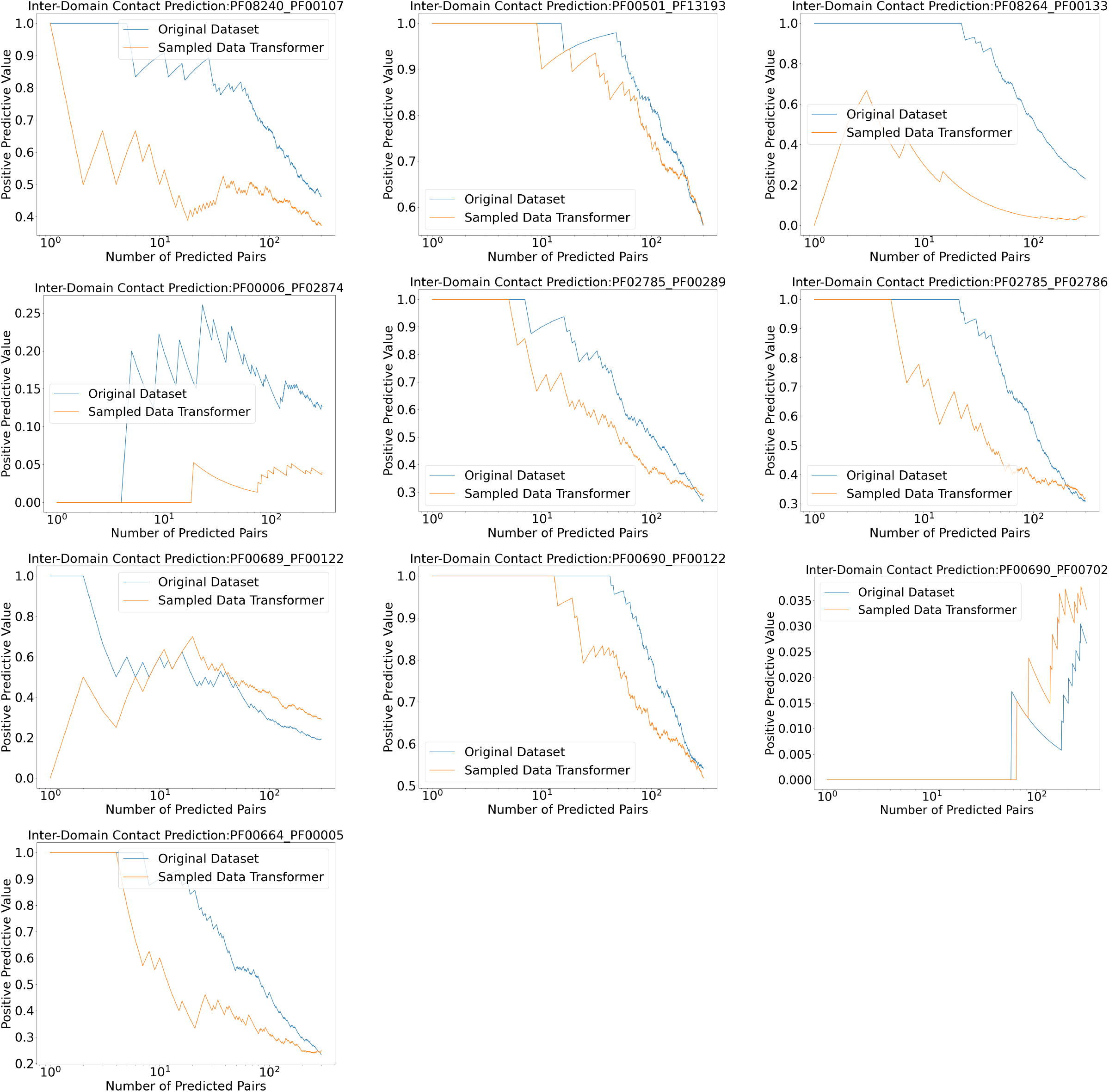
Contact Prediction using plmDCA for the original training set, and a sampled dataset from the the shallow Transformer. The curves represent the Positive Predictive Value (fraction of true positives) with respect to the number of predicted contacts.

### Appendix E Additional Results on Generalization

#### E.1 Matching Performance for Different Distances from Training Set

Here we plot the ratio of correctly matched pairs in the validation set, separated into below-median and above-median distance from the training set.

**Figure E.1:**
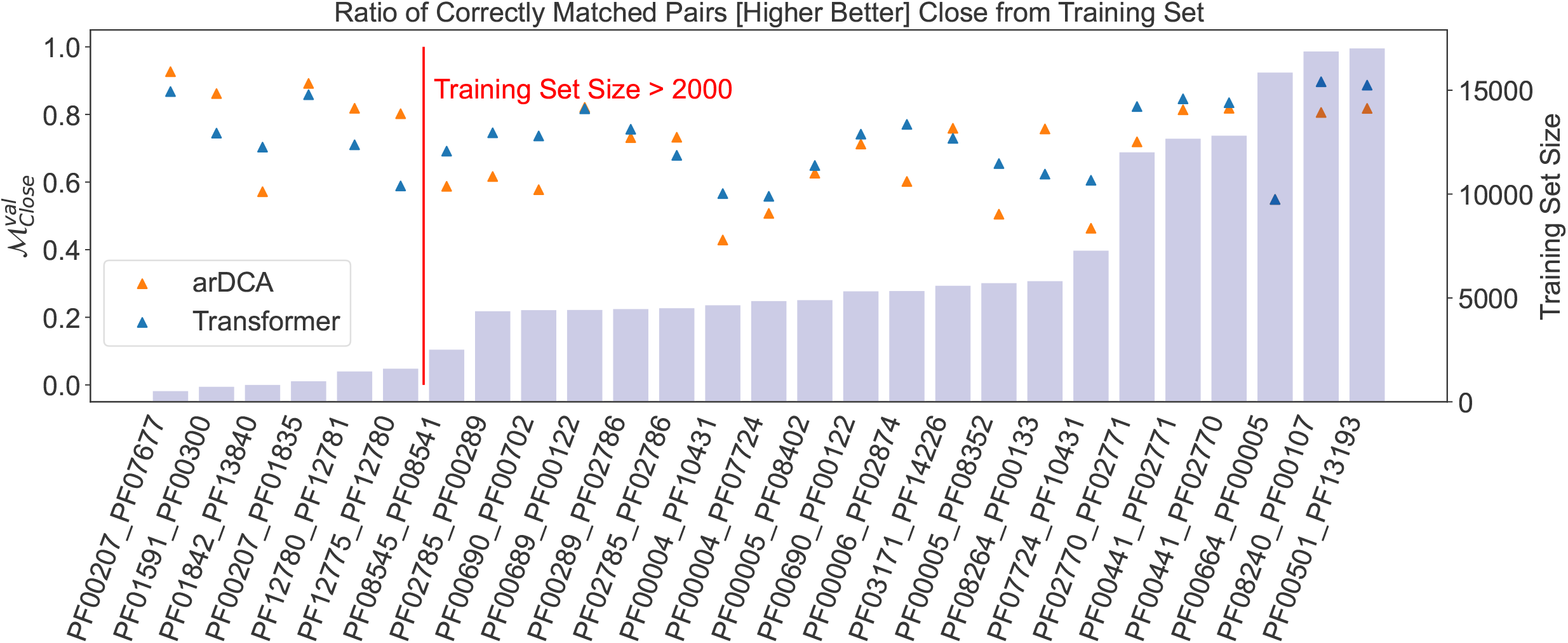
Fraction of correctly matched pairs in the validation set for the shallow Transformer and arDCA. The families are ordered by training set size. Shown are results for the 50% of sequences closest to the training set.

**Figure E.2:**
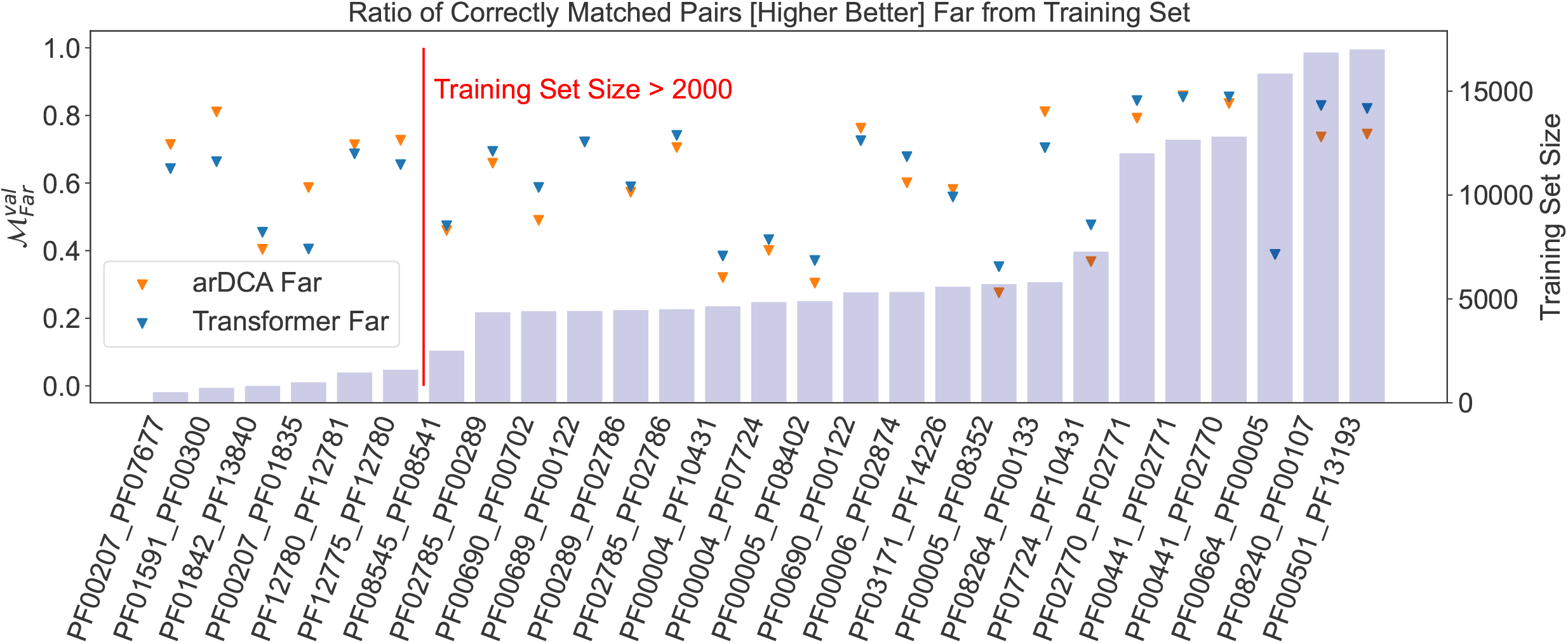
Fraction of correctly matched pairs in the validation set for the shallow Transformer and arDCA. The families are ordered by training set size. Shown are results for the 50% of sequences farthest to the training set.

#### E.2 Accuracy and Perplexity with Distance from Training Set

In this section we present the results for each pair of the analysis of Sec. 4.5.

**Figure E.3:**
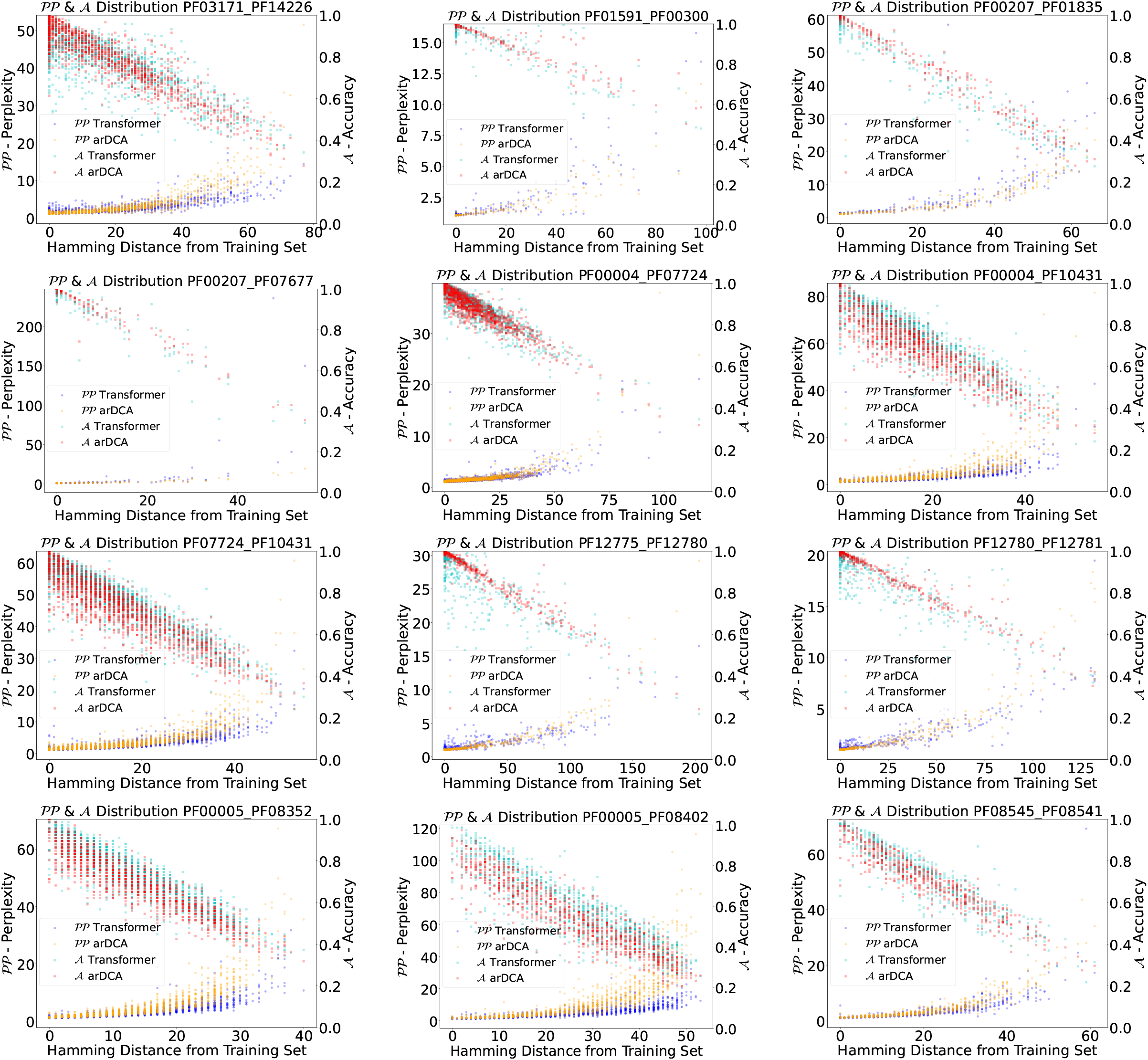
Distribution of the perplexity 𝒫𝒫 and the accuracy 𝒜 of every sequence pair in the validation set with respect to their distance from the training set for the shallow Transformer and arDCA. To fit the page format, we splitted the results on the different families in two figures: this one and the following

**Figure E.4:**
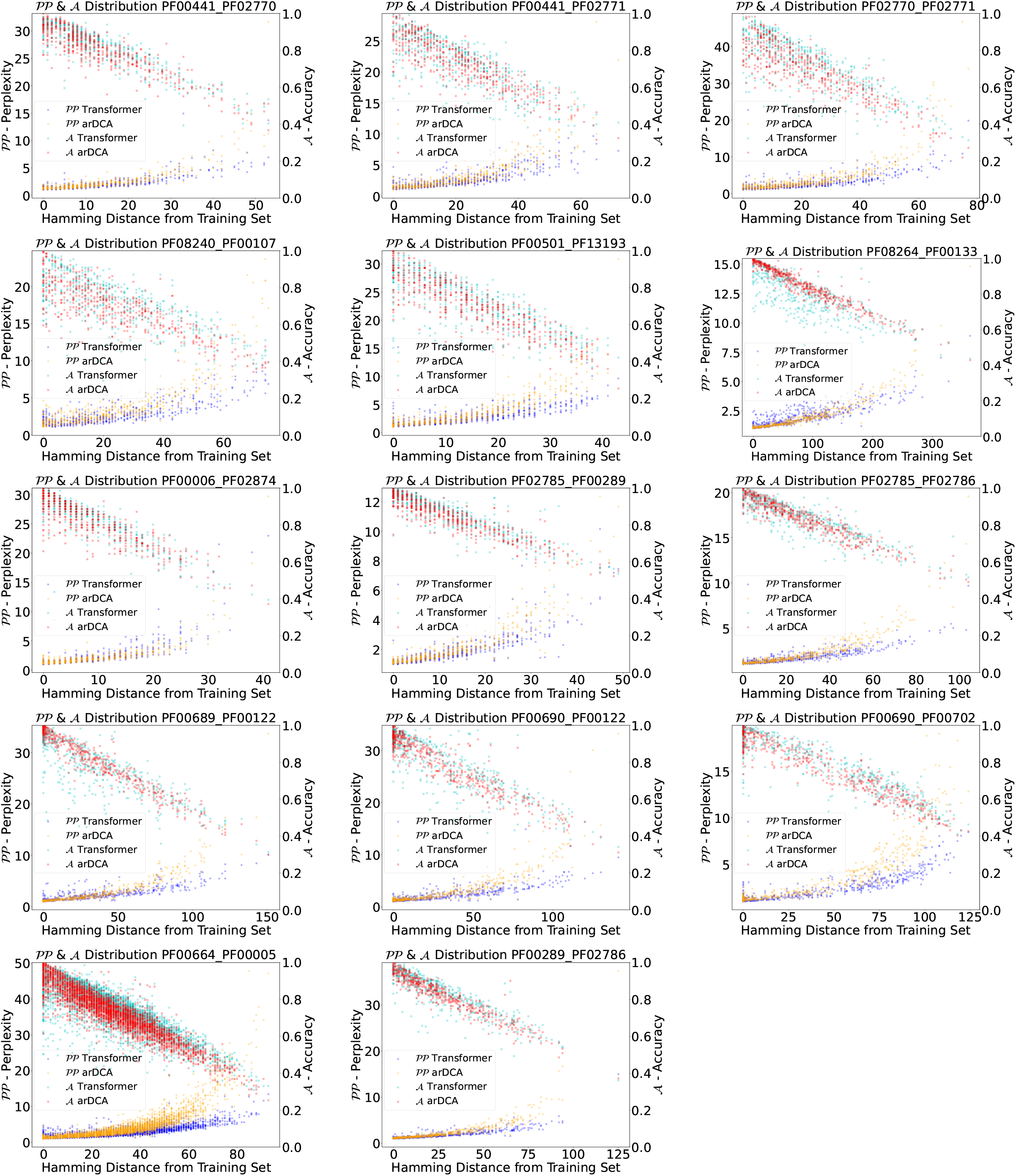
Distribution of the perplexity 𝒫𝒫 and the accuracy 𝒜of every sequence pair in the validation set with respect to their distance from the training set for the shallow Transformer and arDCA.

### Appendix F Additional Matching evaluation

For each protein family we measure the fraction of correctly matched pairs when restricting the problem to the first *n* sequence pairs.

**Figure F.1:**
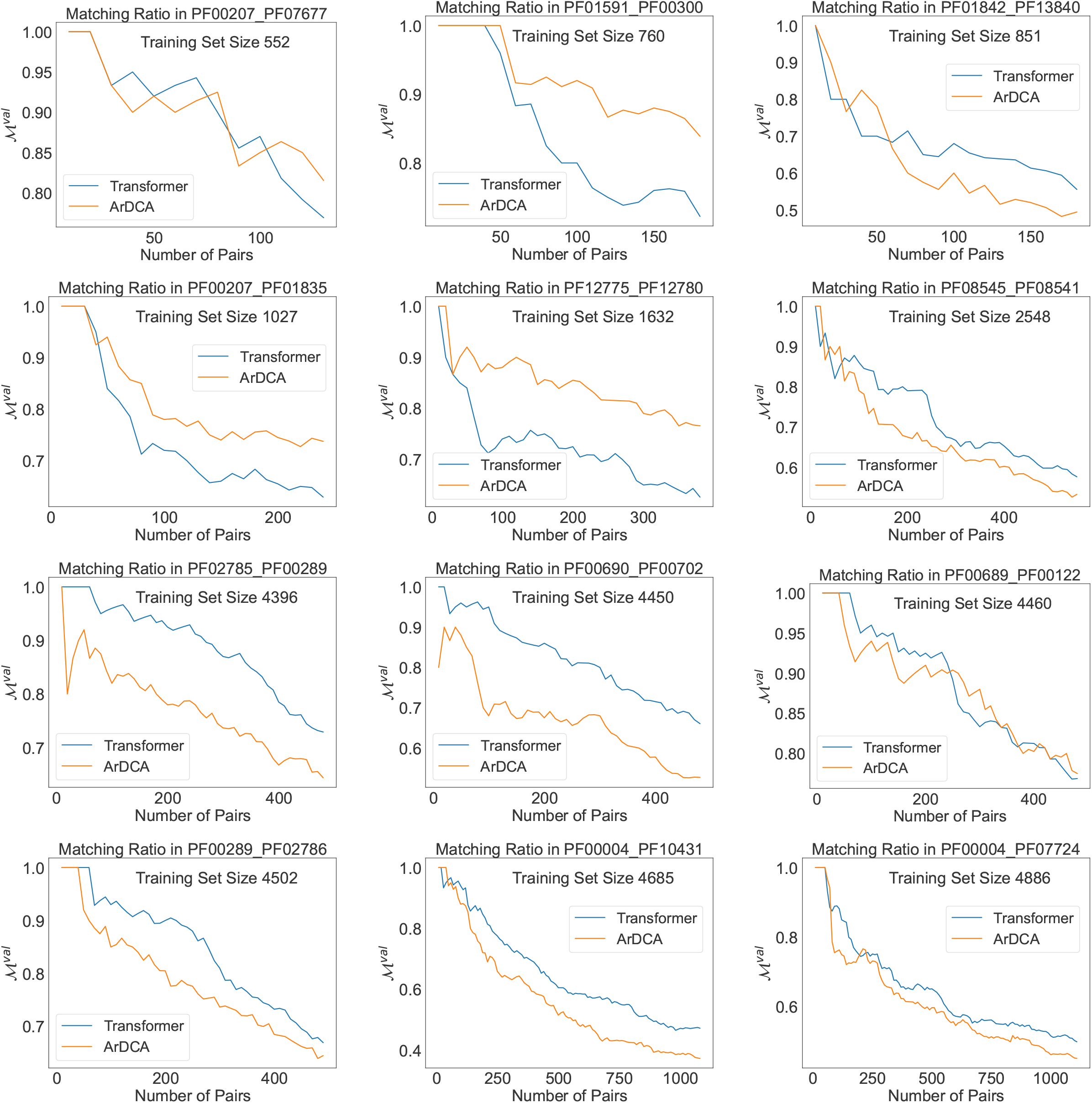
Ratio of correctly matched pairs ℳ^*val*^ for increasing number of pairs for different families. Shown are results for the shallow Transformer (blue) and arDCA (orange). Only a subset of the families are shown in order to save computational resources. The families are ordered according to the training set size. In order to fit the page format, we split the results in this and the next figure.

**Figure F.2:**
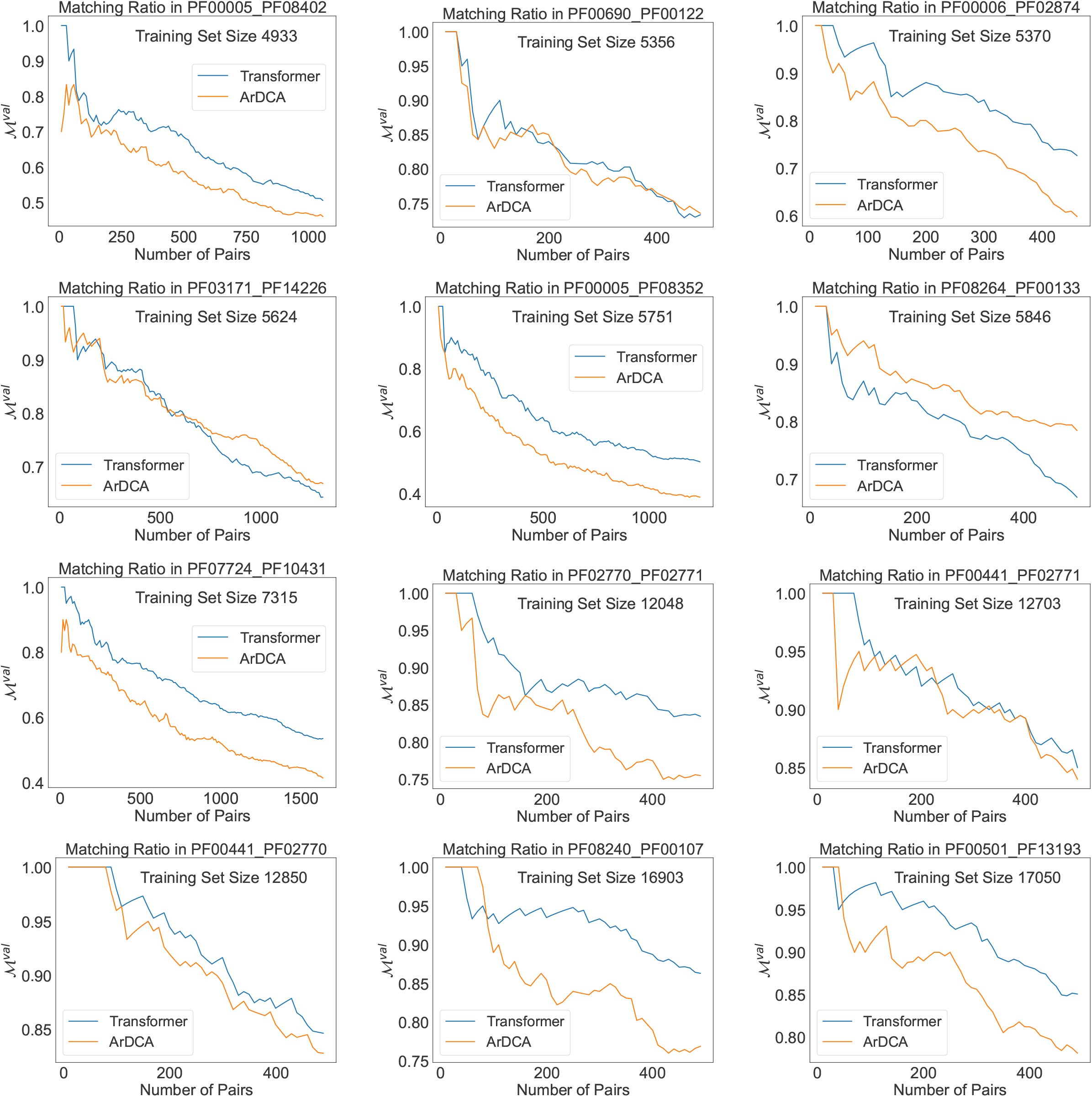
Ratio of correctly matched pairs ℳ^*val*^ for increasing number of pairs for different families. Shown are results for the shallow Transformer (blue) and arDCA (orange). Only a subset of the families are shown in order to save computational resources. The families are ordered according to the training set size. In order to fit the page format, we split the results in this and the previous figure.

#### F.1 D

##### F.1.1 D.3 Sequence Logo and Loss

In Fig. F.3 we show the perplexity per position for family pair PF013171-PF14226, together with the sequence logo. It is evident that biologically conserved positions correspond to lower perplexities, which is to be expected.

**Figure F.3:**
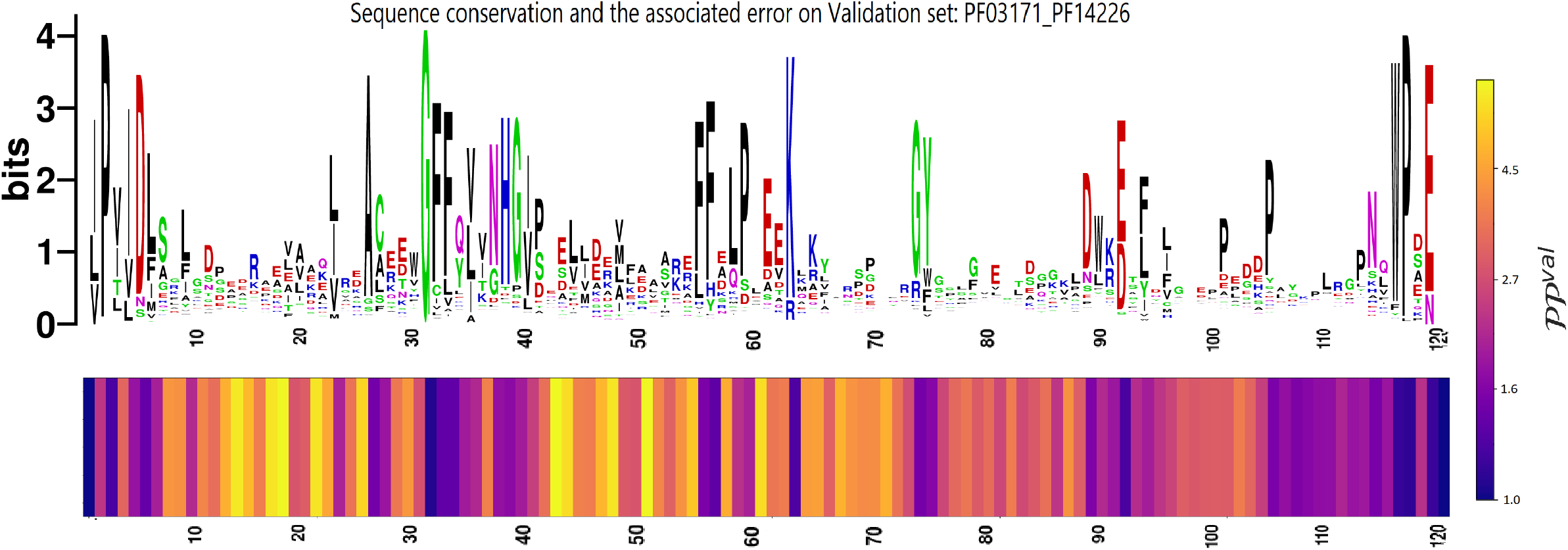
Top: Sequence logo of PF013171-PF14226 paired MSA. Bottom: Distribution of the perplexity with respect to the positions. The errors are concentrated on the most variable position, highlighting that the transformer has understood the basic site wise structure of the distribution.

